# Decoys provide a scalable platform for the genetic analysis of plant E3 ubiquitin ligases that regulate circadian clock function

**DOI:** 10.1101/501965

**Authors:** Ann Feke, Wei Liu, Jing Hong, Man-Wah Li, Chin-Mei Lee, Elton Zhou, Joshua M. Gendron

## Abstract

The circadian clock in all eukaryotes relies on the regulated degradation of clock proteins to maintain 24-hour rhythmicity. Despite this knowledge, we know very few of the components that mediate degradation of proteins to control clock function. This is likely due to high levels of gene duplication and functional redundancy within plant E3 ubiquitin ligase gene families. In order to overcome this issue and discover E3 ubiquitin ligases that control circadian clock function, we generated a library of transgenic Arabidopsis lines expressing dominant-negative “decoy” E3 ubiquitin ligases. We determined their effects on the plant circadian clock and identified dozens of new potential regulators of circadian clock function. To demonstrate the potency of the decoy screening methodology to overcome genetic redundancy and identify *bona fide* clock regulators, we performed follow-up studies on PUB59 and PUB60. Using knock-out studies, we show that they redundantly control circadian clock period by regulating gene splicing. Furthermore, we confirm that they are part of a conserved protein complex that mediates splicing in eukaryotes. This work demonstrates the viability of E3 ubiquitin ligase decoys as a scalable screening platform to overcome traditional genetic challenges and discover E3 ubiquitin ligases that regulate plant developmental processes.

## INTRODUCTION

The circadian clock is essential for proper coordination of biological processes with the environment. In plants, the circadian clock controls diverse aspects of plant development, including hypocotyl elongation, leaf movement, seasonal flowering time, and stress responses, both biotic and abiotic (Dowson-Day and Millar, 1999; Fowler et al., 1999; Hoshizaki and Hamner, 1964; Ingle and Roden, 2014; Liu et al., 2013; Nakamichi et al.,. The timing of the clock is set by environmental inputs, such as daily changes in light and temperature, but it is also self-sustaining and capable of maintaining roughly 24 hour rhythms in the absence of changes in the environmental signals. These self-sustaining oscillations are driven by interlocking transcriptional feedback loops that result in successive expression of a series of transcriptional repressors and activators throughout the day (McClung, 2014; Ronald and Davis, 2017).

In plants, the transcriptional repressors consist predominantly of three groups of proteins. These are the morning expressed CIRCADIAN CLOCK ASSOCIATED 1 (CCA1) and LATE ELONGATED HYPOCOTYL (LHY), the morning and afternoon expressed PSEUDORESPONSE REGULATOR (PRR) family, and the evening expressed evening complex (including EARLY FLOWERING 3(ELF3), EARLY FLOWERING 4 (ELF4) and LUX ARRYTHMO (LUX) (Alabadi et al., 2001; Alabadí et al., 2002; Carré and Kim, 2002; Doyle et al., 2002; Farré et al., 2005; Fujimore et al., 2005; Gendron et al., 2012; Hazen et al., 2005; Helfer et al., 2011; Hicks et al., 1996, 2001; Kikis et al., 2005; Mizoguchi et al., 2002; Nakamichi et al., 2005b, 2005a; Onai and Ishiura, 2005; Schaffer et al., 1998; Wang and Tobin, 1998). More recently, the LIGHT-REGULATED WD (LWD), REVEILLE (RVE), and NIGHT LIGHT-INDUCIBLE AND CLOCK-REGULATED1 (LNK) genes, were identified as critical transcriptional activators in the plant clock and provide a more comprehensive understanding of the transcriptional feedback loops that drive oscillations (Farinas and Mas, 2011; Hsu et al., 2013; Rawat et al., 2011; Wu et al., 2016; Xie et al., 2014).

Eukaryotic circadian clocks employ the ubiquitin proteasome system (UPS) to degrade clock transcription factors at the appropriate time of day (Grima et al., 2002; He et al., 2003; Ito et al., 2012; Ko et al., 2002; Shirogane et al., 2005). The UPS is ideally suited for regulation of the circadian clock because it can mediate protein degradation quickly and specifically. To achieve specificity, the UPS leverages E3 ubiquitin ligase proteins (Chen and Hellmann, 2013; Hua and Vierstra, 2011). E3 ubiquitin ligases act as substrate adaptor proteins by bringing the substrate into proximity of an E2 ubiquitin conjugating enzyme to promote substrate ubiquitylation. Once a poly-ubiquitin chain is added to the substrate, it is sent to the proteasome where it is degraded (Vierstra, 2009). E3 ubiquitin ligases exist in multiple families and contain highly diverse protein recognition domains, allowing them to achieve specificity in the system.

F-box proteins are the substrate adaptor component of a larger E3 ubiquitin ligase complex and are utilized by all eukaryotic circadian clocks (Grima et al., 2002; He et al., 2003; Ito et al., 2012; Ko et al., 2002; Shirogane et al., 2005). The complex, abbreviated SCF, consists of S PHASE KINASE-ASSOCIATED PROTEIN 1 (SKP1), CULLIN, RING-BOX1 (RBX1), and the F-box protein (Bai et al., 1996; Deshaies, 1999; Deshaies and Joazeiro, 2009; Hua and Vierstra, 2011; Lechner et al., 2006). A family of three partially redundant F-box proteins, ZEITLUPE (ZTL), LOV KELCH PROTEIN 2 (LKP2), and FLAVIN-BINDING KELCH REPEAT 1 (FKF1), regulate the circadian clock and flowering time in plants (Imaizumi et al., 2005, 2003; Nelson et al., 2000; Schultz et al., 2001; Somers et al., 2000). ZTL, which has the largest impact on clock function, regulates stability of TOC1, PRR5, and CHE (Fujiwara et al., 2008; Kiba et al., 2007; Lee and Feke et al., 2018; Más et al., 2003). Outside of the ZTL family, some evidence suggests that LHY stability is regulated by the non-F-box RING-type E3 ubiquitin ligase SINAT5 (Park et al., 2010). Since the discovery of these E3 ubiquitin ligases, little progress has been made in identifying additional E3 ubiquitin ligases that participate in clock function.

The inability to identify plant E3 ubiquitin ligases that regulate the circadian clock is likely due to genetic challenges that hamper traditional forward genetic approaches. In Arabidopsis, gene duplication has led to expansion of the genes involved in UPS function (Navarro-Quezada et al., 2013; Risseeuw et al., 2003; Yee and Goring, 2009). For instance, there are approximately 700 Arabidopsis F-box genes, while in humans there are 69 (Finn et al., 2016; Grima et al., 2002; Kuroda et al., 2002; Xu et al., 2009). This has likely led to increased functional redundancy rendering gene knockouts an inefficient method to identify function. To support this, the majority of *ztl* mutant alleles are semi-dominant (Kevei et al., 2006; Martin-Tryon et al., 2006; Somers et al., 2004, 2000). This suggests that reverse genetic strategies may be a more potent approach to identify E3 ubiquitin ligases that regulate clock function.

In order to overcome redundancy in plant E3 ubiquitin ligase families, we developed a “decoy” E3 ubiquitin ligase approach. The decoy approach involves expressing an E3 ubiquitin ligase that lacks the ability to recruit the E2 conjugating enzyme but retains the ability to bind to the substrate (Han et al., 2004; Kishi and Yamao, 1998; Latres et al., 1999; Li et al., 2012; Zhou et al., 2015). We have shown that this inactivates the full-length E3 ubiquitin ligase and acts to stabilize the substrate protein (Lee and Feke et al., 2018). The decoy acts as a dominant-negative, making it an effective genetic tool to identify the function of redundant E3 ubiquitin ligases. Additionally, the decoy stabilizes interaction with substrate proteins. This allows us to express the decoy with an affinity tag to study interactions between E3 ubiquitin ligases and substrates.

Here, we demonstrate the potency and scalability of the decoy technique by performing a reverse genetic screen to identify regulators of the circadian clock. We attempted to create decoy-expressing transgenic lines for half of the F-box-type E3 ubiquitin ligases and all of the U-box-type E3 ubiquitin ligases from Arabidopsis. Our completed library contains nearly ¼ of the Arabidopsis E3 ubiquitin ligases (Vierstra, 2009), spanning sixteen different protein-protein interaction domain classes, and including many genes with known functions as well as many that have not been studied in detail previously.

We used the decoy library to identify E3 ubiquitin ligases that can regulate the plant circadian clock. We uncovered a surprisingly large number of genes that regulate clock function with minor effects and a smaller number with more dramatic effects on clock period or phase. We then perform focused genetic studies on *PLANT U-BOX 59* and *PLANT U-BOX 60 (PUB59* and *PUB60*), two homologous U-box genes which have been previously implicated in splicing. We go on to determine their molecular function in the clock by showing that the core clock gene, *PRR9*, is mis-spliced in the *pub59/pub60* double mutant. This work demonstrates the effectiveness of the decoy technique as a screening platform and identifies the first U-box-type E3 ligases that are involved in clock function in any system. It also establishes two important community resources: a list of E3 ligases that regulate the plant circadian clock, and a decoy library that is freely available and can be used to identify E3 ubiquitin ligases involved in any plant developmental processes.

## RESULTS

### Construction of the decoy library

In order to discover E3 ubiquitin ligases that regulate the plant circadian clock, we created a library of transgenic plants expressing decoy E3 ubiquitin ligases. Decoy E3 ubiquitin ligases are identical to the native E3 ubiquitin ligases but lack the domain that recruits the E2 conjugating enzymes. Thus, the decoys retain substrate binding abilities but lack the ability to mediate substrate ubiquitylation (Han et al., 2004; Kishi and Yamao, 1998; Latres et al., 1999; Lee et al., 2018; Zhou et al., 2015). Thus, transgenic plants expressing decoy ubiquitin ligases should act dominantly to endogenous E3 ubiquitin ligases.

In this pilot screen, we started with the F-box family of E3 ubiquitin ligases, in part because of the known role of F-box proteins on the circadian clock and in part because we have demonstrated the viability of the technique with three F-box E3 ubiquitin ligases (Lee and Feke et al., 2018). We selected roughly half of the F-box gene family, including representatives from all of the large classes and most of the small classes of F-box proteins for this initial screen. Some genes that we chose have differential expression under various growth conditions, while there was nothing known about the expression of others.

The F-box domain is unusual in that it is almost always located in the N-terminal portion of the protein. Thus, F-box decoy constructs were created by amplifying the sequence downstream of the F-box domain in each gene and creating pENTR vectors for each. The decoy constructs were then recombined into a vector that will drive their expression under a *CaMV (Cauliflower Mosaic Virus) 35S* promoter and in-frame with a 6xHis/3xFLAG affinity tag (Fig 1a).

**Figure 1.**
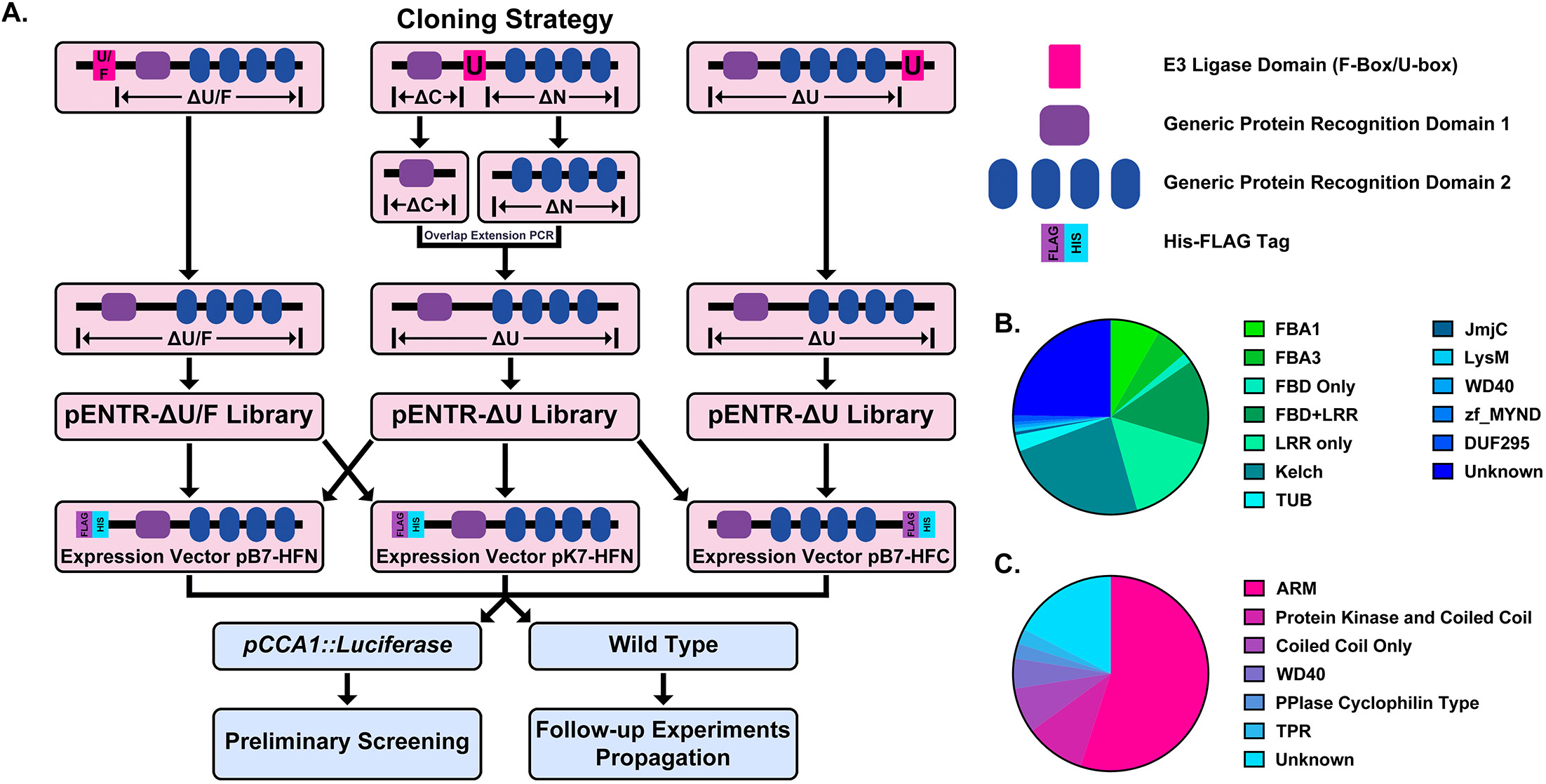
Construction of the Arabidopsis Decoy Library. A) Cloning and experimental workflow of the F-box and U-box decoy library. F-box decoys follow the same path as the N-terminal U-box decoys. B) Distribution of protein recognition domains in the F-box decoy library C) Distribution of protein recognition domains in the U-box decoy library.

In order to test the decoy technique’s viability across E3 ubiquitin ligase classes, we also selected a second family of E3 ubiquitin ligases to include in the library. The U-box family was selected due to its small size, containing only around 60 members (Azevedo et al., 2001; Finn et al., 2016; Yee and Goring, 2009). Furthermore, the U-box domain itself is well-defined and roughly the same size as the F-box domain (Andersen et al., 2004; Aravind and Koonin, 2000). Unlike the F-box domain, which is characteristically in the N-terminus of the protein, the U-box domain can be located anywhere throughout the protein sequence. For U-boxes with the U-box domain in the C– or the N-terminus, we amplified all sequence that was located upstream or downstream of the U-box, respectively. For those with the U-box domain located in the middle, we amplified both upstream and downstream sequences and then ligated the two halves together, a successful strategy that we utilized in our previous study (Lee and Feke et al., 2018). The decoy constructs were then recombined into the same expression vectors as described for the F-box decoy library (Fig 1a).

Based on another large scale cloning project in Arabidopsis, we expected 70-80% success rate in cloning the F-box genes (Pruneda-Paz et al., 2014). In fact, we were able to clone 82% of the attempted F-box genes, and ultimately succeeded in isolating transgenic plants for 65% of cloned decoy F-boxes (Table S1). The inability to isolate transgenic plants for the remaining 35% of cloned decoy F-boxes may be due to a multitude of factors, including but not limited to lethality caused by expressing the decoy, reduced transformation efficiency, or other technical constraints. Of those successfully generated transgenic lines, the majority contained either an LRR, Kelch, or F-box Associated (FBA1, FBA3, or FBD) protein recognition domain (at 30%, 24%, or 30%, respectively), with some F-box proteins containing both LRR and FBD domains together (14%) (Fig 1b). The remaining F-boxes contained a small number of other domains (6%), including TUBBY-like or WD40 domains, or no known protein recognition domain (25%). We also generated transgenic plants expressing 65% of the U-box family (Table S1). Of those cloned, 55% contained ARM repeats, 10% contained a Protein Kinase domain, 7.5% contained only a coiled coil region, 5% contained a WD repeat, 5% contained other annotated domains, and 17.5% contained no annotated domains (Fig 1c).

In sum, we attempted to generate a transgenic library expressing decoys for approximately 1/4^th^ of the Arabidopsis E3 ubiquitin ligases (Vierstra, 2009). From here on, we use the term “decoy” to describe a transgenic plant containing the *35S* promoter driven, FLAG-His tagged E3 ubiquitin ligase with the E3 ubiquitin ligase domain deleted. A decoy “line” is defined as a single, independent T1 insertion line containing a decoy construct, and a decoy “population” is a group of decoy lines which all express the same decoy transgene but are independent T1 transgenics.

### Screen design

In order to identify E3 ubiquitin ligases that regulate clock function, we transformed our decoy library into transgenic Col-0 plants harboring the *CIRCADIAN CLOCK ASSOCIATED* 1 promoter driving the expression of the *Luciferase* gene (*CCAlp::Ludferase*) and monitored clock function (Pruneda-Paz et al., 2009). From automated imaging experiments performed under constant light conditions on week-old seedlings entrained in LD (12 hours light/12 hours dark) conditions, we were able to measure clock period, phase, and relative amplitude error (RAE – a statistical measure of rhythmicity (Moore et al., 2014; Zielinski et al., 2014)) of all transgenic lines and controls.

As a quality control measure, we first filtered our data for those with reliable control experiments. For an experiment to be included in the analyses we required that the control *CCAlp::Lucifernse* populations have a standard deviation of less than 0.75 hours. We chose this threshold value as it equates to the closest 15 minute window to a 95% confidence interval within a 24 hour period. We removed any experiments with larger control variances from further analyses. By discarding datasets with larger degrees of variation, we reduce the chances of false positives and reduce the impact of any unpredictable environmental differences.

### The role of E3 ubiquitin ligase decoys in clock rhythmicity

Some circadian clock mutants completely ablate clock function and cause arrhythmicity (Hazen et al., 2005; Nakamichi et al., 2009). To determine the rhythmicity of the decoy lines we calculated the RAE for all 8502 individual T1 transgenic lines. We plotted each F-box and U-box gene from the screen on trees so that any potential redundant genes would be nearer to each other (Dereeper et al., 2008). The trees do not have evolutionary significance and only provide relative gene relatedness at the protein sequence level and were created using the full-length rather than decoy sequence. Five individual lines had an RAE greater than 0.6 which signifies lack of rhythmicity (Fig S1–S2). In comparison, no control lines (n=1783) had an RAE greater than 0.6 (Fig S3a, S4a). No decoy populations had more than one arrhythmic line, making it unlikely that any decoy ablates clock function. Rather, it is possible that the insertion landed in a gene necessary for rhythmicity in these lines. The lack of arrhythmicity in decoy lines is not surprising, and supports prior studies that show post-translational degradation mechanisms are often not necessary for rhythmicity (Hurley et al., 2016; Larrondo et al., 2015; Ode et al., 2017)

### The effects of E3 ubiquitin ligase decoys on clock phase

We next determined whether the decoy populations have alterations in phasing of the Arabidopsis circadian clock. We calculated phase difference for each transgenic line. This was done by calculating the average phase of the control population in each experiment, then subtracting this value from the phase value of each individual T1 transgenic line analyzed in the same experiment. Individual control lines were normalized in this same manner (Fig S3b, S4b). Interestingly, we observed a phase shift in most of our decoy expressing populations when compared to the wild type, with the large majority showing a significant phase advance (Fig 2–3). This approximately 1 hour advance in phase (compare to Fig S3b, S4b) appeared to be a general effect of transgene expression in our experiments, suggesting the phase of the *CCAlp::Luciferase* is particularly sensitive to transgene overexpression.

**Figure 2.**
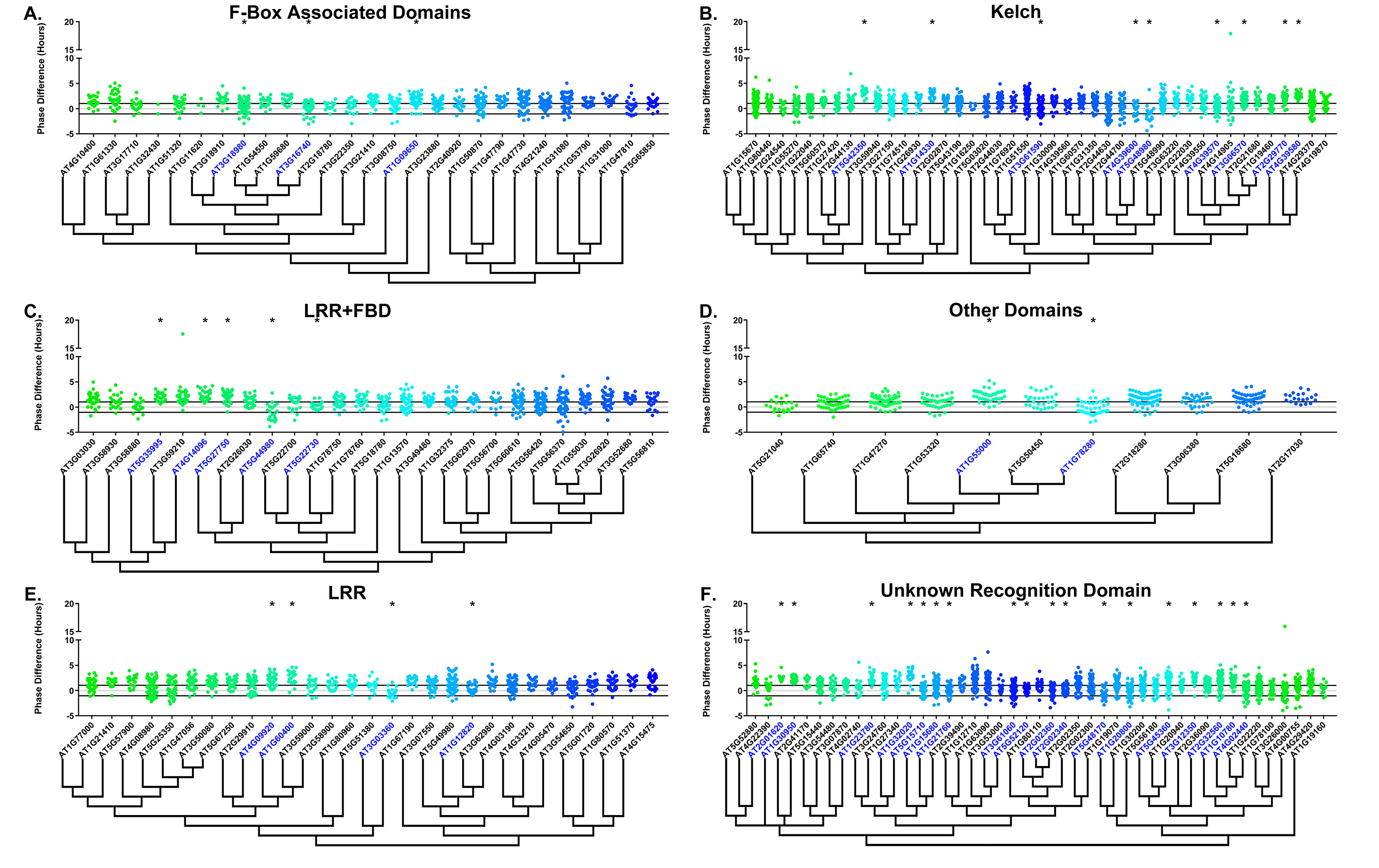
Phase distributions of F-box decoy lines. Values presented are the difference between the period of the individual decoy line and the average period of the *CCA1p::Luciferase* control in the accompanying experiment. The grey line is at the average control value and the black lines are at +/– the standard deviation of the control lines. Genes are separated by protein recognition domain and ordered by closest protein homology using Phylogeny.Fr, (Dereeper et al., 2008), and a tree showing that homology is displayed beneath the graph. F-Box Associated Domains = FBA1, FBA3, and FBD only. Other Domains = TUB, JmjC, LysM, WD40, zf_MYND, and DUF295. * and blue gene names = The entire population differs from wildtype with a Bonferroni-corrected p <1.94×10^-4^. Only data from those experiments where the control *pCCA1::Luciferaselines* display a standard deviation less than 0.75 were included in our analyses.

**Figure 3.**
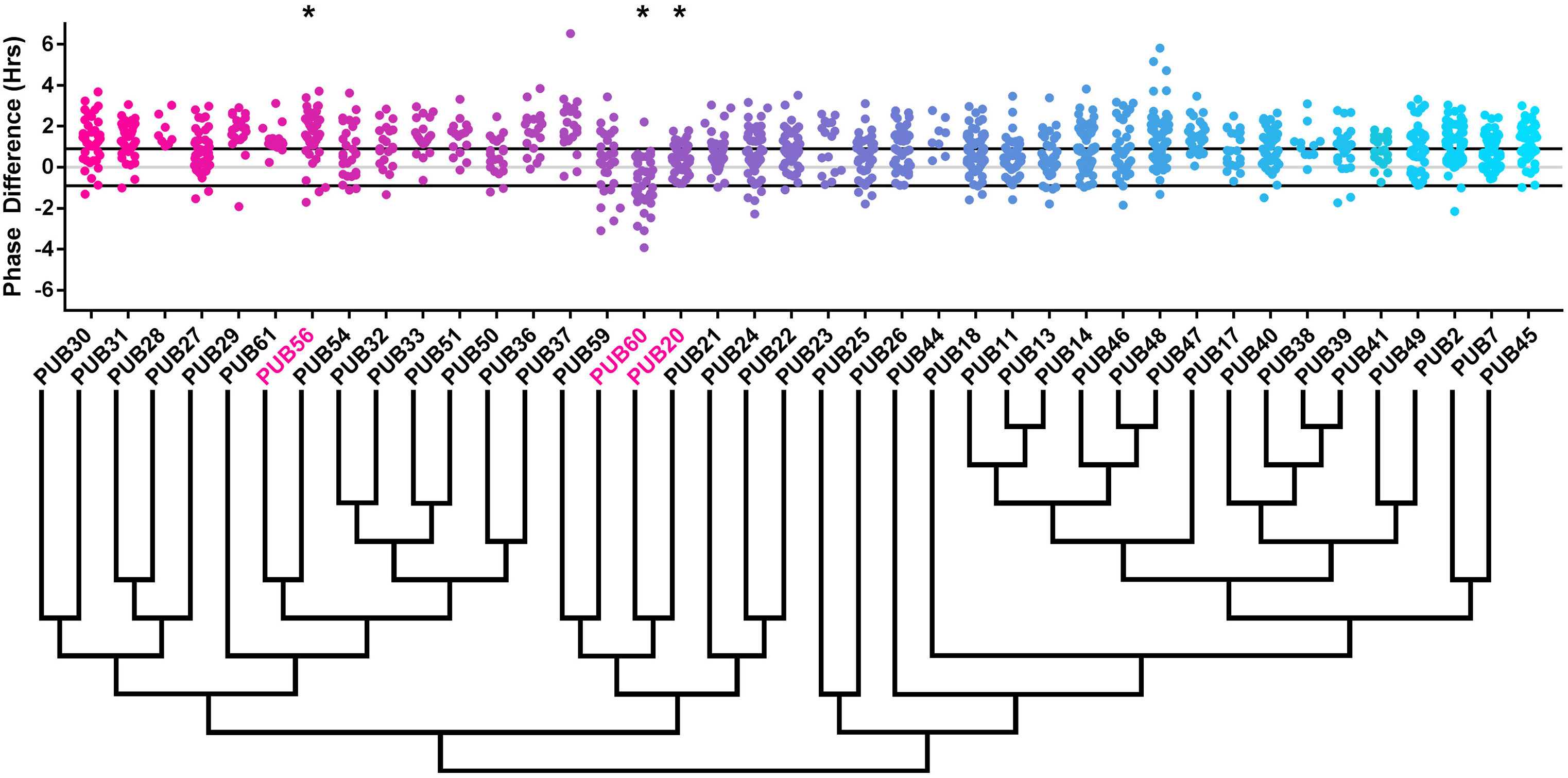
Phase distributions of U-box decoy lines. Values presented are the difference between the period of the individual decoy line and the average period of the *CCA1p::Luciferase* control in the accompanying experiment. The grey line is at the average control value and the black lines are at +/– the standard deviation of the control lines. Genes are ordered by closest protein homology using Phylogeny.Fr, (Dereeper et al., 2008), and a tree showing that homology is displayed beneath the graph. * and pink gene names = The entire population differs from wildtype with a Bonferroni-corrected p <1.00×10^-3^.

To overcome the general phase advance we compared each decoy population to the entire set of decoy populations for statistical testing. Using this method, we found that 40 F-box decoy populations cause a statistically significant change in phase (Welch’s t-test with a Bonferroni corrected α of 1.94×10^-4^), equating to approximately 22% of tested populationss (Fig 2, marked with stars and blue gene names). We next set a cutoff of two hours phase difference to further subdivide our group into major (>2 hours) and minor (<2 hours) regulators. Based on this definition, many of the F-box decoy populations had minor phase differences (37/40 populations) while only three (*AT5G48980* – 2.36 hours delayed, *AT5G44980* – 2.16 hours delayed, and *AT5G42350* – 2.16 hours advanced) had major phase differences. In addition to the F-box decoys, three of the U-box decoy populations have phase differences (Welch’s t-test with a Bonferroni corrected α of 1.00×10^-3^), none of which had major phase differences (Fig 3).

### Sequence and Expression Analysis of Phase-Regulating F-box Proteins

Two of the three major phase regulators have not been studied previously. For this reason, we propose to name them *ALTERED CLOCK F-BOX 1(ACF1 – AT5G44980*) and *ACF2 (AT5G48980)*. As *AT5G42350* is already known as *COP9 INTERACTING F-BOXKELCH1 (CFK1)*, we do not give this gene the *ACF* nomenclature.

In order to understand the function and regulation of the *ACFs*, we mined publically available expression data and the literature. ACF1, which contains both an F-box associated domain (FBD) and Leucine rich repeats (LRR), has no publications detailing its function. The absence of an identified phenotype may be due to the existence of a close homolog (*AT5G44960* – E-value of 1.60×10^-146^, Table 1), as higher order mutants or dominant negative technology, such as the decoy technique, may be required to uncover its function.

**Table 1.**
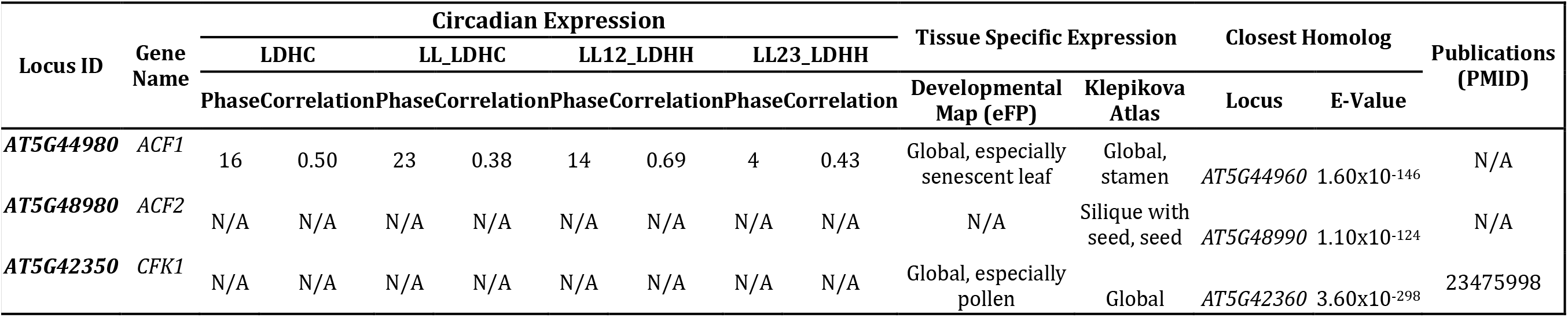
Publically available data for strong candidate F-box phase regulators. Circadian expression data is from the Diurnal Project gene expression tool (Mockler et al., 2007). Tissue specific expression is from the Arabidopsis eFP browser (Klepikova et al., 2016; Winter et al., 2007). The closest homolog was determined by WU-BLAST2 using BLASTP and the Araport11 protein sequences database. N/A indicates that data is not available.

| Locus ID | Gene Name | Circadian Expression |  |  |  |  |  |  |  | Tissue Specific Expression |  | Closest Homolog |  | Publications (PMID) |
| --- | --- | --- | --- | --- | --- | --- | --- | --- | --- | --- | --- | --- | --- | --- |
|  |  | LDHC |  | LL_LDHC |  | LL12_LDHH |  | LL23_LDHH |  |  |  | Locus | E-Value |  |
|  |  | Phase | Correlation | Phase | Correlation | Phase | Correlation | Phase | Correlation | Developmental Map (eFP) | Klepikova Atlas |  |  |  |
| AT5G44980 | ACF1 | 16 | 0.50 | 23 | 0.38 | 14 | 0.69 | 4 | 0.43 | Global, especially senescent leaf | Global, stamen | AT5G44960 | 1.60x10<sup>-146</sup> | N/A |
| AT5G48980 | ACF2 | N/A | N/A | N/A | N/A | N/A | N/A | N/A | N/A | N/A | Silique with seed, seed | AT5G48990 | 1.10x10<sup>-124</sup> | N/A |
| AT5G42350 | CFK1 | N/A | N/A | N/A | N/A | N/A | N/A | N/A | N/A | Global, especially pollen | Global | AT5G42360 | 3.60x10<sup>-298</sup> | 23475998 |

Often genes that regulate clock function are also themselves regulated by the clock or light. Thus, we attempted to determine whether *ACF1* is regulated by the circadian clock or diurnal cycles. *ACF1* is not regulated by either light cycles or the circadian clock, as the correlation value, a measure of the similarity between the expression data and the hypothesized cycling pattern, is less than the standard correlation cutoff of 0.8 (Mockler et al., 2007). Furthermore, many core clock genes are expressed ubiquitously in the plant, although tissue specific clocks do exist (Endo et al., 2014; Lee and Seo, 2018; Shimizu et al., 2015). For this reason, we searched two expression atlases to determine the expression patterns of *ACF1*. Tissue expression maps suggest that *ACF1* is expressed globally, although there may be some enrichment in senescent leaves or anthers (Klepikova et al., 2016; Winter et al., 2007). The global expression pattern suggests that *ACF2* has the potential to be involved in phasing of the circadian clock in all plant tissues.

We performed the same analysis on *ACF2*, which contains a Kelch repeat. No publications are available, possibly due to the existence of a close homolog (*AT5G48990* – E-value of 1.10×10^-124^, Table 1), although *ACF2* was described in a manuscript discussing the prevalence of the Kelch-repeat containing F-box proteins in Arabidopsis (Andrade et al., 2001). Diurnal and circadian expression data was not available for this gene (Mockler et al., 2007). While data on *ACF2* expression is unavailable in one expression maps, the second shows predominant expression in seed (Klepikova et al., 2016; Winter et al., 2007).

More is known about *CFK1*. While temporal expression data is unavailable, expression maps demonstrate that *CFK1* is expressed globally (Table 1) (Klepikova et al., 2016; Mockler et al., 2007; Winter et al., 2007), suggesting it is not tissue specific. *cfk1* knockout mutants cause decreased hypocotyl length, and interact with the CONSTITUTIVE PHOTOMORPHOGENIC 9 (COP9) Signalosome (Franciosini et al., 2013). *CFK1* has a very close homolog (*AT5G42360*, also known as *CFK2* – E-value of 3.6×10^-298^), but while it cannot be completely redundant with *CFK1* because of the knockout phenotype, reduction in the levels of both genes increases the phenotypic severity (Franciosini et al., 2013). *CFK1* is expression induced by light (Franciosini et al., 2013), providing strength to the argument that it could be involved in clock function.

### The role of F-box decoys in clock period

Many clock E3 ubiquitin ligases control periodicity, thus we determined whether the decoys cause changes in clock period (Godinho et al., 2007; Han et al., 2004; Liu et al., 2018; Reischl et al., 2007). We calculated the period difference by calculating the average period of the control population in each experiment, then subtracting this value from the period value of each individual T1 transgenic line analyzed in the same experiment. Unlike the effects on phase, we did not observe a general period shift across all decoy lines when doing this analysis (Fig 4, compare to Fig S3c). This suggests that the period of the *pCCA1::Luciferase* reporter is not sensitive to general effects of transgene overexpression. From this analysis we found that 36 F-box decoy populations have significantly different periods than the control (Welch’s t-test with a Bonferroni corrected α of 2.55×10^-4^) (Fig 4, marked with stars and green gene names). These correspond to approximately 19% of tested populations (Fig 4, marked with stars and green gene names). We next divided the group into populations with minor (<1 hours period difference) and major (>1 hour period difference) effects on the period difference. Interestingly, only one F-box decoy population has a major period difference (*AT1G20800* – 1.02 hours longer than the wild type). The remaining have minor effects on the period difference.

**Figure 4.**
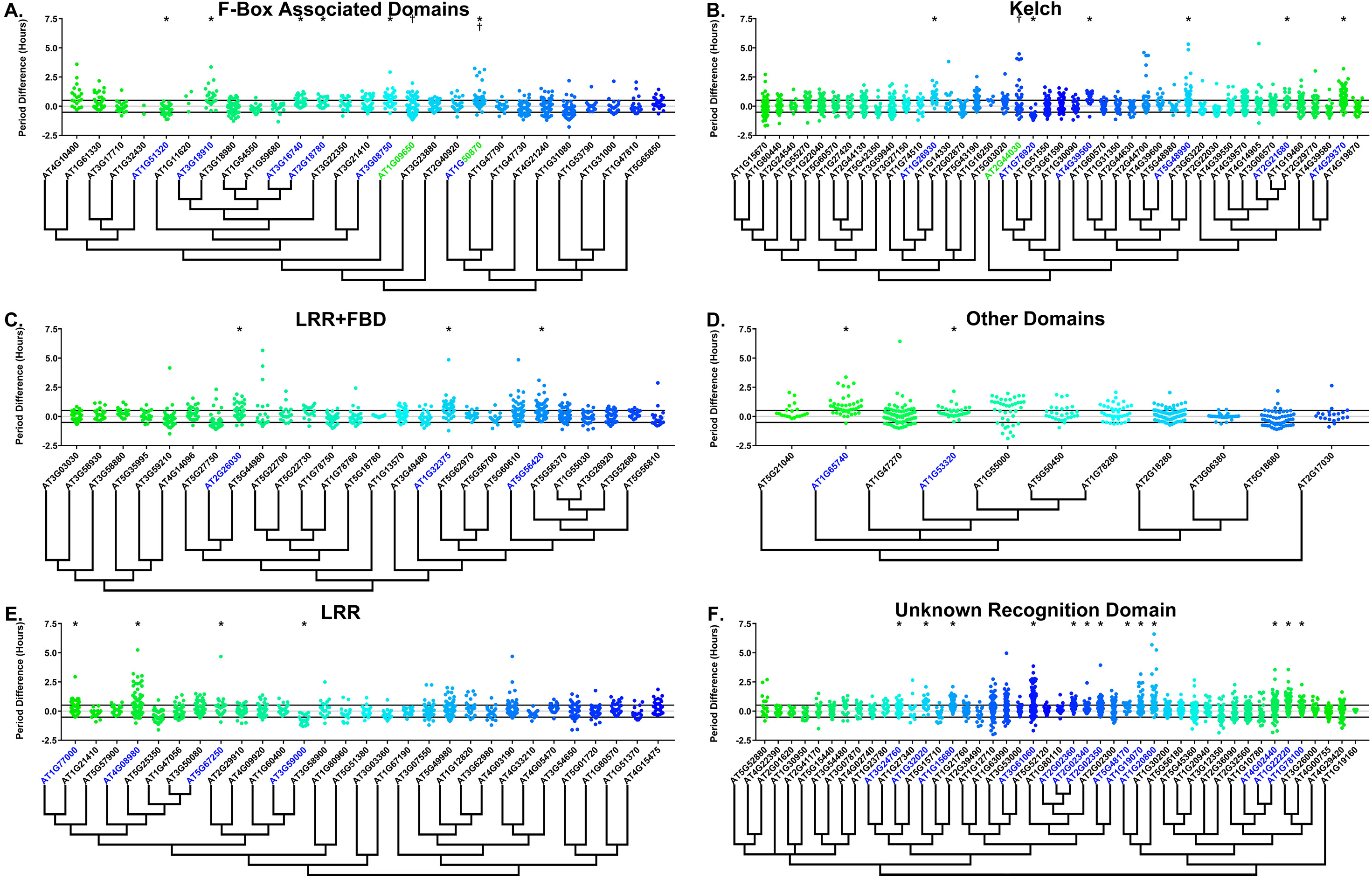
Period distributions of F-box decoy lines. Values presented are the difference between the period of the individual decoy line and the average period of the *CCA1p::Luciferase* control in the accompanying experiment. The grey line is at the average control value and the black lines are at +/– the standard deviation of the control lines. Genes are separated by protein recognition domain and ordered by closest protein homology using Phylogeny.Fr, (Dereeper et al., 2008), and a tree showing that homology is displayed beneath the graph. F-Box Associated Domains = FBA1, FBA3, and FBD only. Other Domains = TUB, JmjC, LysM, WD40, zf_MYND, and DUF295. * and blue gene names = The entire population differs from wildtype with a Bonferroni-corrected p <2.55×10^-4^; f and green gene names = A subset of the population differs from wildtype with a Bonferroni-corrected p <2.55×10^-4^. Only data from those experiments where the control *CCA1p::Luciferase* lines display a standard deviation less than 0.75 were included in our analyses.

Previously we showed that expressing decoys of clock-regulating F-box genes can result in separable subpopulations that affect circadian period differentially (Lee and Feke et al., 2018). We further analyzed the period data to identify decoy populations with statistically separable subpopulations. We define a subpopulation as a group of three or more decoy lines that have similar periods to each other but are statistically different than other subpopulations from the same decoy population (see the materials and methods section for further details). There are two F-box decoy populations that are not different than the control as a whole population, but have distinct subpopulations that are different than the control (Fig 4, marked with daggers and blue gene names). One, *AT2G44030*, has a subpopulation that has a major effect on period, containing four lines with an average period 4.1 hours longer than the control. The remaining subpopulation is not significantly different from the control. A second decoy population, *AT1G50870*, which has a minor effect on the period overall (0.57 hours longer) also contains two separable subpopulations. One subpopulation has a major effect on the period (2.8 hours longer, n=5), while the second has a minor effect (0.36 hours longer, n=55).

Only three of the 41 identified populations or subpopulations had shorter periods than the wildtype. *AT1G76920* and *AT1G51320* both have shorter periods overall (0.76 and 0.29 hours shorter, respectively), while *AT1G09650* has a subpopulation that is 0.61 hours shorter than the wildtype. Nothing to date has been published regarding the functions of any of these genes. While none of these falls into the defined major effect category, the relative scarcity of short period effects make these potential candidates for follow-up study.

As a quality control measure we examine Luciferase reporter traces to identify any abnormalities in the rhythms of the decoy populations. We plotted the average Luciferase traces for the F-box decoy populations (or those with subpopulations) with major effects on period difference (Fig 5). Additionally, we have plotted the raw period data, color coded by subpopulation, so that the lines being included in each of the traces are obvious. The traces show that the decoys have relatively normal rhythms aside from the shifts in period, suggesting that the decoys are affecting period but not phasing and rhythmicity.

**Figure 5.**
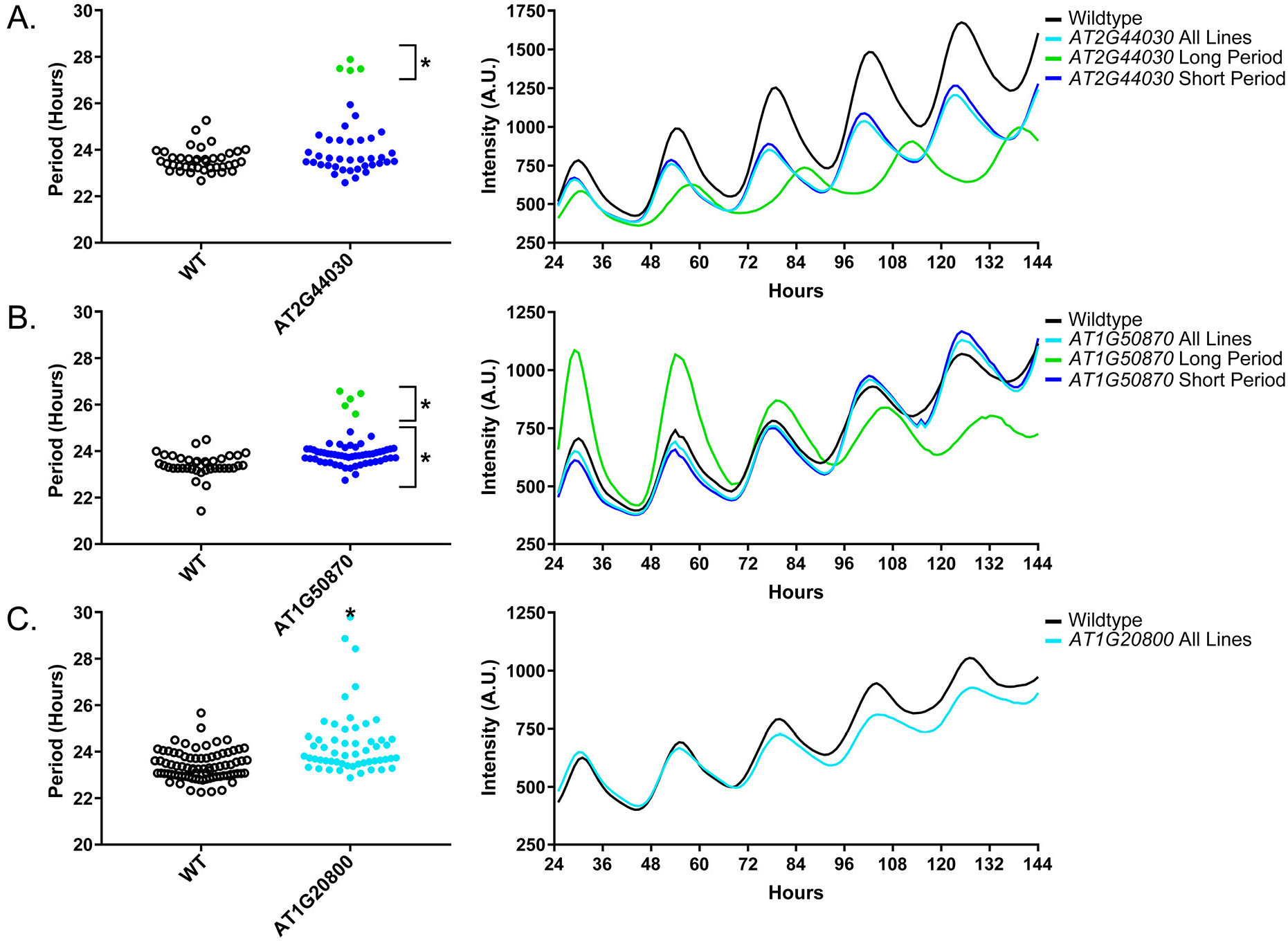
Circadian Phenotypes for selected high-priority F-box decoy lines. Period values and average traces for decoy lines with significant differences from the control across the entire population or a sub-population of lines greater than 1. Period values presented are raw period lengths as determined by *CCA1p::Luciferase*, and traces are calculated from the average image intensity across each seedling at each hour throughout the duration of the imaging experiment. Time 0 is defined as the dawn of the release into LL. A) *AT2g44030* decoy. B) *AT1G50870* decoy. C) *AT1G20800* decoy. Brackets define individual groups used for statistical testing against the wildtype control using a Welch’s t-test with a Bonferroni-corrected α of 2.55×10^-4^. * represents p<α. When multiple subpopulations were detected, the members of each group were separately averaged and presented in the traces along with the average of all lines.

### Sequence and Expression Analysis of Period-Regulating F-box Proteins

We also give the *ACF* nomenclature to the three genes with major period differences. *AT1G20800* we name *ACF3, AT2G44030* we name *ACF4*, and *AT1G50870* we name *ACF5*. We also performed the expression analyses and literature searches on these three *ACF* genes.

No publications exist regarding the functions of ACF3, which also has no known protein recognition domain. *ACF3* contains a close homolog (*AT1G20803* – E-value of 1.6×10^-87^] which may be why it was not identified in previous forward genetic screens (Table 2). Expression of *ACF3* is not controlled by the circadian clock or diurnal cycles (Mockler et al., 2007). The two expression maps have widely varying expression data for *ACF3*, as one suggests that this gene is expressed globally, while the other suggests that it is expressed exclusively in floral buds (Klepikova et al., 2016; Winter et al., 2007).

**Table 2.** Publically available data for strong candidate F-box period regulators. Circadian expression data is from the Diurnal Project gene expression tool (Mockler et al., 2007). Tissue specific expression is from the Arabidopsis eFP browser (Klepikova et al., 2016; Winter et al., 2007). The closest homolog was determined by WU-BLAST2 using BLASTP and the Araport11 protein sequences database. N/A indicates that data is not available.

| Locus ID | Gene Name | Circadian Expression |  |  |  |  |  |  |  | Tissue Specific Expression |  | Closest Homolog |  | Publications (PMID) |
| --- | --- | --- | --- | --- | --- | --- | --- | --- | --- | --- | --- | --- | --- | --- |
|  |  | LDHC |  | LL_LDHC |  | LL12_LDHH |  | LL23_LDHH |  | Developmental Map (eFP) | Klepikova Atlas | Locus | E-Value |  |
|  |  | Phase | Correlation | Phase | Correlation | Phase | Correlation | Phase | Correlation |  |  |  |  |  |
| AT1G20800 | ACF3 | 18 | 0.51 | 4 | 0.66 | 15 | 0.65 | 16 | 0.66 | N/A | Young flower, stamen, ovule | AT1G20803 | 1.6x10<sup>-87</sup> | N/A |
| AT2G44030 | ACF4 | 18 | 0.73 | 5 | 0.54 | 6 | 0.62 | 22 | 0.79 | Seed, pollen | Young stamen, flower buds | AT3G46050 | 2.5x10<sup>-59</sup> | N/A |
| AT1G50870 | ACF5 | N/A | N/A | N/A | N/A | N/A | N/A | N/A | N/A | N/A |  | AT1G47790 | 1.2x10<sup>-102</sup> | N/A |

ACF4, which contains a Kelch-repeat domain, also has no described mutant phenotype, and is only described in an overview of the Kelch-repeat containing F-box proteins in Arabidopsis (Andrade et al., 2001). The closest homolog to *ACF4 (AT3G46050* – E-value of 2.5×10^-59^) has not been studied (Table 2). As with *ACF3, ACF4* is not controlled by the circadian clock or light cycles (Mockler et al., 2007). *ACF4* is expressed globally in one developmental map and in floral buds in the other (Klepikova et al., 2016; Winter et al., 2007).

Very little is known about *ACF5* outside of its predicted FBA3 domain. No expression data, either diurnal or circadian regulation or tissue-specific, is available for *ACF5* (Table 2). Furthermore, no studies have been published on the function of this gene. It has some homology to other genes (*AT1G47790* – E-value of 1.2×10^-102^), which again indicates that it may have been missed in previous genetic screens due to redundancy.

### The role of U-box decoys in clock period

We also analyzed the U-box library data to identify decoy populations or subpopulations with period differences. We found that two populations of decoy lines had average period lengths longer than wild type, *PUB60* (1.31 hours longer) and *PUB41* (0.37 hours longer) (Welch’s t-test with a Bonferroni corrected α of 1.09×10^-3^) (Fig 6, marked with stars and pink gene names) making *PUB60* the only U-box decoy with a major effect on the average period difference.

**Figure 6.**
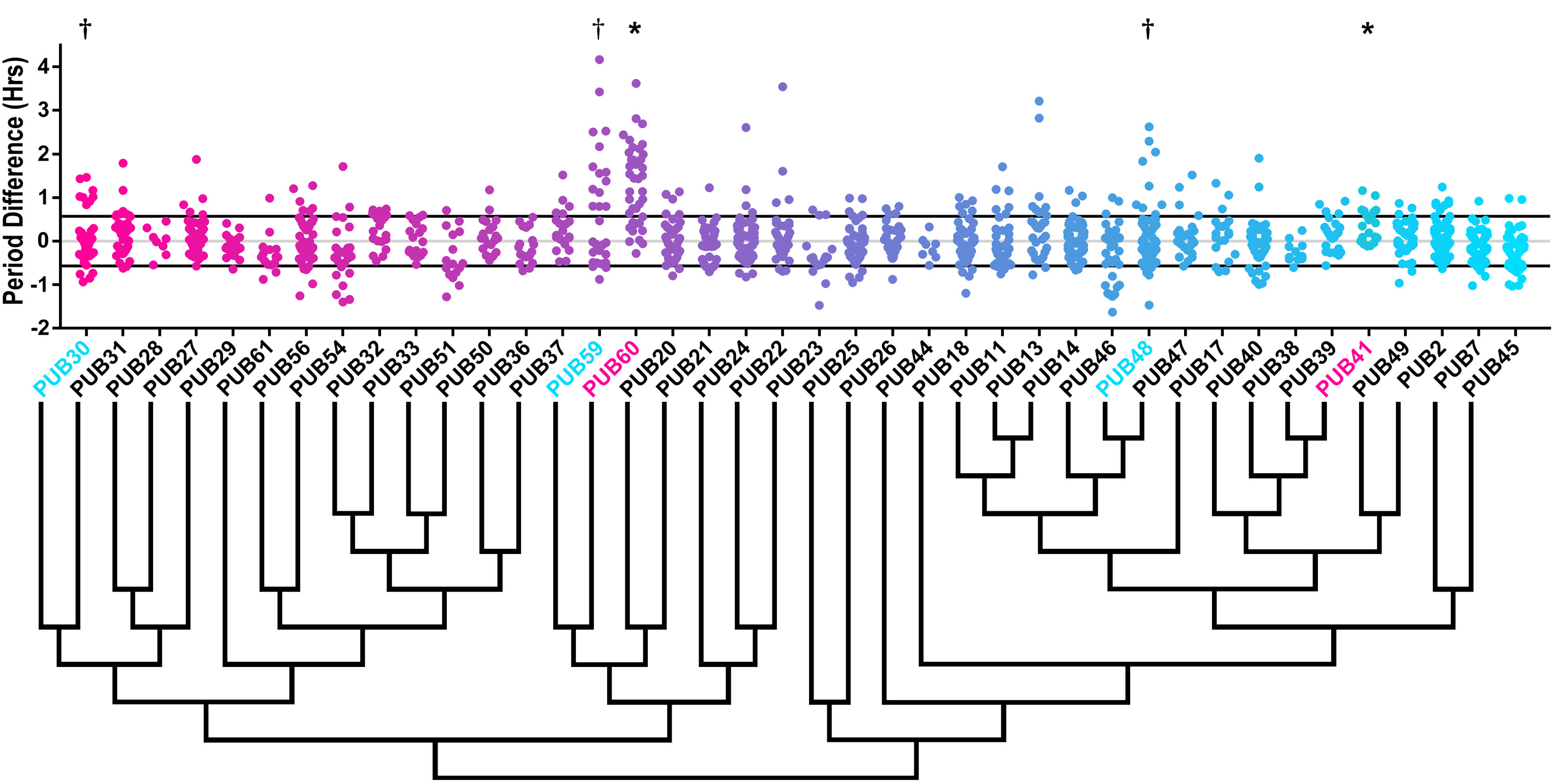
Period distributions of U-box decoy lines. Values presented are the difference between the period of the individual decoy line and the average period of the *CCA1p::Luciferase* control in the accompanying experiment. The grey line is at the average control value and the black lines are at +/– the standard deviation of the control lines. Genes are ordered by closest protein homology using Phylogeny.Fr, (Dereeper et al., 2008), and a tree showing that homology is displayed beneath the graph. * and pink gene names = The entire population differs from wildtype with a Bonferroni-corrected p <1.09×10^-3^; † and cyan gene names = A subset of the population differs from wildtype with a Bonferroni-corrected p <1.09×10^-3^.

Three additional U-box decoy populations had subpopulations that were different from the control, *PUB30, PUB48*, and *PUB59* (Fig 6, marked with daggers and blue gene names). In this case all three had subpopulations that we consider strong regulators. *PUB30* has one subpopulation which is 1.11 hours longer than the wildtype (n=7), while the other subpopulation is not significantly different (n=32). Similarly, *PUB48* has one subpopulation which is 2.20 hours longer than the wildtype (n=4), and a second subpopulation which is not significantly different (n=48). *PUB59* is different in that it has one subpopulation that we consider in the major effect category (1.75 hours longer, n=15), and one that is short period (0.4 hours shorter, n=14). Because these subpopulations alter the period differently, the overall average is not statistically significant from wildtype (Fig 6).

Again we examined Luciferase reporter traces to identify any abnormalities in the rhythms of the decoy populations. We plotted the average Luciferase traces for the U-box decoy populations (or those with subpopulations) with major effects on period difference (Fig 7). Additionally, we have plotted the raw period data, color coded by subpopulation, so that the lines being included in the traces are obvious. *PUB30, PUB59*, and *PUB60* decoy populations or subpopulations have period defects but otherwise normal rhythms. *PUB48* long period lines, on the other hand, show some rhythmic abnormality. The traces appear to decrease in rhythmicity on day four (Fig 7b) but then regain rhythmicity on day six. Interestingly, some of the control lines for this experiment have abnormally long periods suggesting some stochastic noise in this experiment (Fig 7b). Although this subpopulation and the controls pass our stringent statistical filters, the trace and raw period data may suggest that the results need to be interpreted carefully for *PUB48* and further experimentation will be required to confirm its role in the clock.

**Figure 7.**
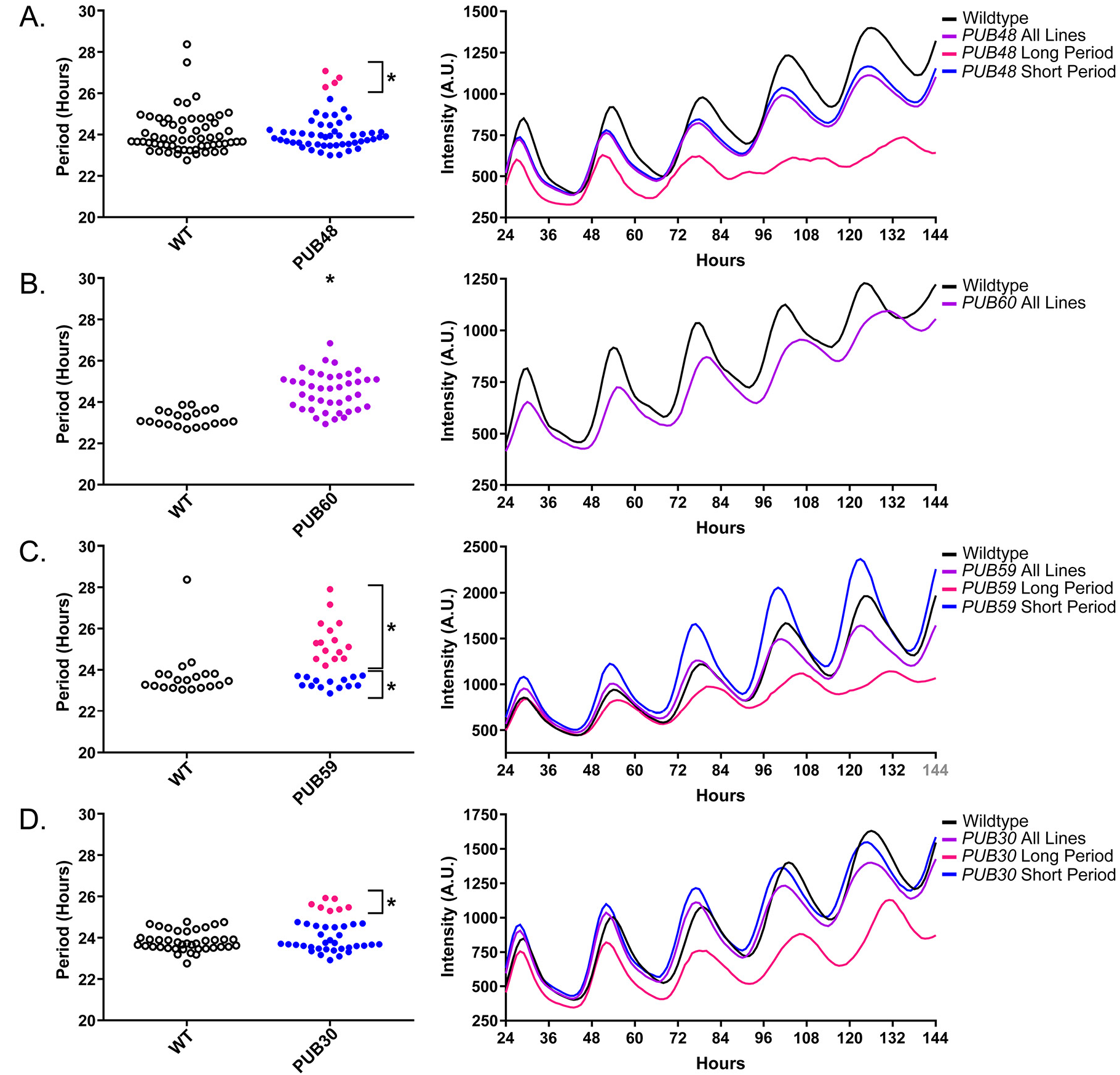
Circadian Phenotypes for selected high-priority U-box decoy lines. Period values and average traces for decoy lines with significant differences from the control across the entire population or a sub-population of lines greater than 1. Period values presented are raw period lengths as determined by *CCA1p::Luciferase*, and traces are calculated from the average image intensity across each seedling at each hour throughout the duration of the imaging experiment. Time 0 is defined as the dawn of the release into LL. A) *PUB48* decoy. B) *PUB59* decoy. C) *PUB60* decoy. D) *PUB30* decoy. Brackets define individual groups used for statistical testing against the wildtype control using a Welch’s t-test with a Bonferroni-corrected α of 1.09×10^-3^. * represents p<α. When multiple subpopulations were detected, the members of each group were separately averaged and presented in the traces along with the average of all lines.

### Sequence and Expression Analysis of Period-Regulating U-box Proteins

As the U-box genes have been given their PUB nomenclature (Azevedo et al., 2001; Yee and Goring, 2009), we did not rename these genes. We do, however, perform the same expression and literature searches that we performed on the *ACF* genes. *PUB30*, which contains two armadillo (ARM) repeats, has a potential homolog, *PUB31* (E-value of 1.2×10^-171^, Table 3). *PUB30* expression is rhythmic under diurnal conditions and under one of the circadian conditions (Mockler et al., 2007). Both tissue expression maps suggest that *PUB30* is expressed globally (Klepikova et al., 2016; Winter et al., 2007). *PUB30* has a described function in inhibiting the salt stress response (Hwang et al., 2015; Zhang et al., 2017).

**Table 3.**
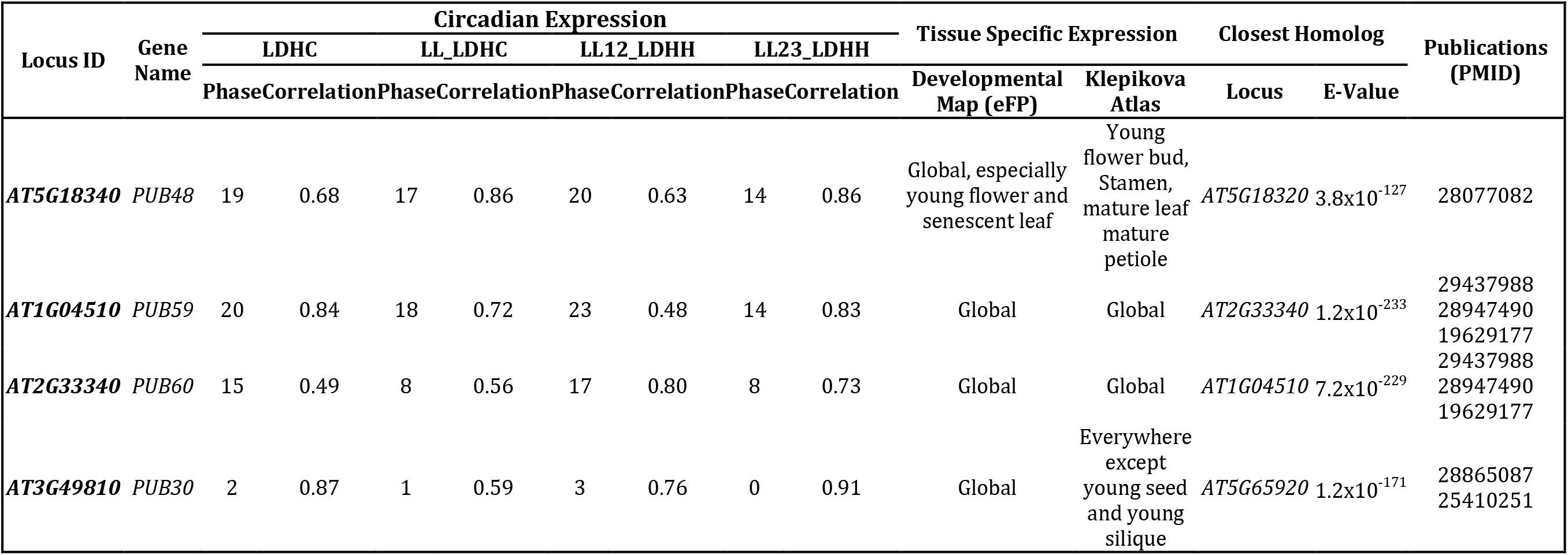
Publically available data for strong candidate U-box period regulators. Circadian expression data is from the Diurnal Project gene expression tool (Mockler et al., 2007). Tissue specific expression is from the Arabidopsis eFP browser (Klepikova et al., 2016; Winter et al., 2007). The closest homolog was determined by WU-BLAST2 using BLASTP and the Araport11 protein sequences database.

| Locus ID | Gene Name | Circadian Expression |  |  |  |  |  |  |  | Tissue Specific Expression |  | Closest Homolog |  | Publications (PMID) |
| --- | --- | --- | --- | --- | --- | --- | --- | --- | --- | --- | --- | --- | --- | --- |
|  |  | LDHC |  | LL_LDHC |  | LL12_LDHH |  | LL23_LDHH |  |  |  | Locus | E-Value |  |
|  |  | Phase | Correlation | Phase | Correlation | Phase | Correlation | Phase | Correlation | Developmental Map (eFP) | Klepikova Atlas |  |  |  |
| AT5G18340 | PUB48 | 19 | 0.68 | 17 | 0.86 | 20 | 0.63 | 14 | 0.86 | Global, especially young flower and senescent leaf | Young flower bud, Stamen, mature leaf, mature petiole | AT5G18320 | 3.8x10<sup>-127</sup> | 28077082 |
| AT1G04510 | PUB59 | 20 | 0.84 | 18 | 0.72 | 23 | 0.48 | 14 | 0.83 | Global | Global | AT2G33340 | 1.2x10<sup>-233</sup> | 29437988<br>28947490<br>19629177 |
| AT2G33340 | PUB60 | 15 | 0.49 | 8 | 0.56 | 17 | 0.80 | 8 | 0.73 | Global | Global | AT1G04510 | 7.2x10<sup>-229</sup> | 29437988<br>28947490<br>19629177 |
| AT3G49810 | PUB30 | 2 | 0.87 | 1 | 0.59 | 3 | 0.76 | 0 | 0.91 | Global | Everywhere except young seed and young silique | AT5G65920 | 1.2x10<sup>-171</sup> | 28865087<br>25410251 |

*PUB48*, like *PUB30*, contains two ARM repeats and has a partially redundant homolog, *PUB46* (Table 3)(Adler et al., 2017). *PUB48* expression is not rhythmic under diurnal cycles. Interestingly, it is rhythmic under circadian conditions in two out of three available experiments, and has a similar phase in both (ZT14 and ZT17). Expression profiling suggests that *PUB48* is expressed globally, although it exhibits some enrichment in floral buds and potentially senescent leaves (Klepikova et al., 2016; Winter et al., 2007). *PUB48* is also involved in stress responses, as it has been shown to positively regulate the response to drought stress (Adler et al., 2017).

*PUB59* and *PUB60* both contain 7 WD repeats, are 82% identical at the protein sequence level, and are known to act redundantly (Table 3) (Monaghan et al., 2009). While both cycle under one of the circadian conditions, only *PUB59* cycles under diurnal conditions (Mockler et al., 2007). Tissue-specific expression profiles are very similar for the two genes, as both are expressed globally (Klepikova et al., 2016; Winter et al., 2007). *PUB59* and *PUB60* are orthologous to the human and yeast Pre-mRNA Processing factor 19 (PRP19) proteins, the central components of the spliceosome activation machinery known as the Nineteen Complex (NTC) (Chanarat and Sträßer, 2013; Hogg et al., 2010). In plants, they were initially identified regulators of plant immunity, but further work confirmed their roles in the regulation of splicing and miRNA biogenesis (Jia et al., 2017; Li et al., 2018; Monaghan et al., 2009). Accordingly, splicing is a critical regulatory step in plant clock function (Filichkin and Mockler, 2012; Park et al., 2012; Sanchez et al., 2010; Simpson et al., 2016; Wang et al., 2012). Because the *PUB59* and *PUB60* decoys affect period, we hypothesize that they are redundantly controlling clock function through regulated splicing of clock genes.

### PUB59 and PUB60 are functionally redundant U-box proteins that regulate plant circadian clock function

To prove the validity of our decoy screening platform we performed detailed genetic and molecular follow-up experiments on *PUB59* and *PUB60*, two potentially redundant regulators of the plant circadian clock. We grew the *pub59* and *pub60* single knockout mutants (Monaghan et al., 2009) in LD conditions for 10 days and transferred them to constant light for two days. We collected tissue from the plants every three hours for two days and performed qRT-PCR to measure expression of the core clock genes, *CCA1* and *TOC1*. The *pub59* and *pub60* single mutants alone have little effect on the period, amplitude, or phase of the circadian clock (Fig 8A-D) suggesting their functions may be redundant. Thus, we obtained the *pub59/pub60* double mutant and monitored *CCA1* and *TOC1* expression under the same conditions as the single mutants (Monaghan et al., 2009). Unlike the single mutants, the *pub59/pub60* double mutant has a significant phase delay (Fig 8E-F). We quantified the phases using FFT-NLLS analysis on the Biodare2 platform (biodare2.ed.ac.uk Zielinski et al., 2014). The *pub59/pub60* double mutant has a phase delay of 3.95 hours for *TOC1* expression and 5.90 hours for *CCA1* expression (Fig 8G). The phase delay is consistent with the effects caused by lengthened clock period, similar to the period lengthening observed in the *PUB59* and *PUB60* decoy populations. This suggests that *PUB59* and *PUB60* are *bona fide* regulators of the circadian clock, and the first U-box genes, to our knowledge, identified as regulators of the circadian clock in any system.

Furthermore, the absence of a clock phenotype in the *pub59* and *pub60* single mutants demonstrates the genes are redundant, and highlights the strength of the decoy technique to overcome traditional genetic barriers.

**Figure 8.**
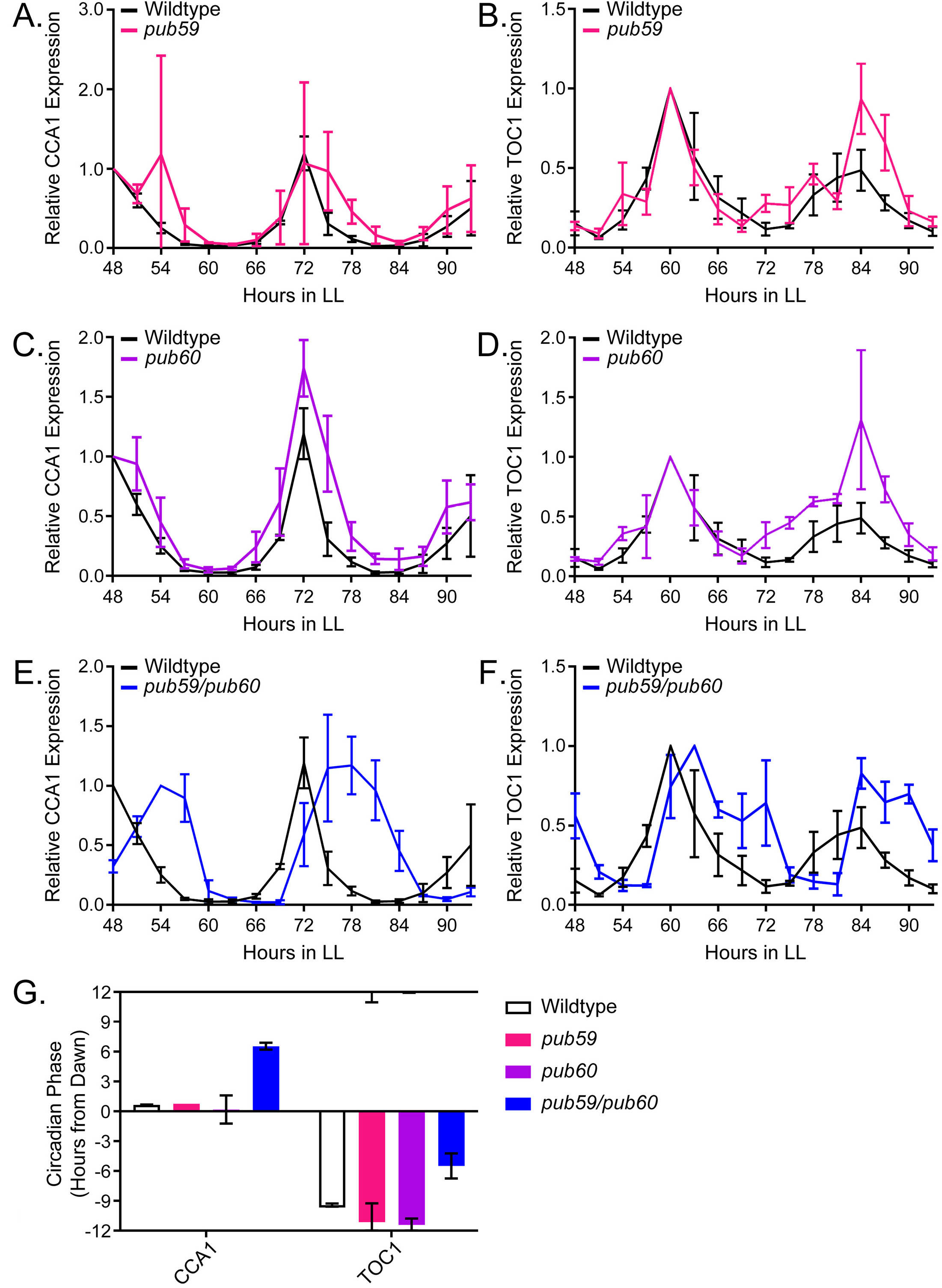
qRT-PCR of clock gene expression in *pub59/pub60* mutant lines. A,C,E) *CCA1* and B,D,F) *TOC1* expression was measured using quantitative RT-PCR in wildtype or homozygous A-B)*pub59*, C-D)*pub60*, and E-F)*pub59/pub60* mutant lines under constant light conditions. Quantifications are the average of three biological replicates with error bars showing standard deviation. G) FFT-NLLS analysis through the Biodare2 analysis platform shows altered phasing in the *pub59/pub60* double mutant. Error bars represent the standard deviation.

### PUB59 and PUB60 control splicing of plant circadian clock genes

As the *pub59/pub60* mutant was recently shown to exhibit global splicing defects and intron retention, we hypothesized that these splicing defects may impact the circadian clock. Like *PUB59* and *PUB60*, another component of the NTC, *SNW/SKI-interacting protein (SKIP*), has a lengthened circadian period when mutated, likely due to the dysregulation of *PRR9* and *PRR7* splicing (Wang et al., 2012). For these reasons, we performed time course qRT-PCR on the *pub59/pub60* mutant to investigate *PRR9* splicing. In the *pub59/pub60* mutant we observe a decrease in the amplitude of the active *PRR9* spliceoform, termed *PRR9a*, as well a secondary inactive form, referred to as *PRR9b* (Fig 9A-B). This is accompanied by an increase in the amplitude of the inactive spliceoform, *PRR9c*, which retains an intron inappropriately (Fig 9C). We quantified these differences in amplitude using the Biodare2 analysis platform (biodare2.ed.ac.uk Zielinski et al., 2014), and observed a similar trend, although the large error bars in the wildtype do overlap with the mutant for *PRR9a* and *PRR9b* (Fig 9D). These results are consistent with previous results showing that mutations in splicing factors result in elevated *PRR9c* expression. Together, this data suggests that *PUB59* and *PUB60* play a role in the circadian clock by promoting the proper splicing of clock components.

**Figure 9.**
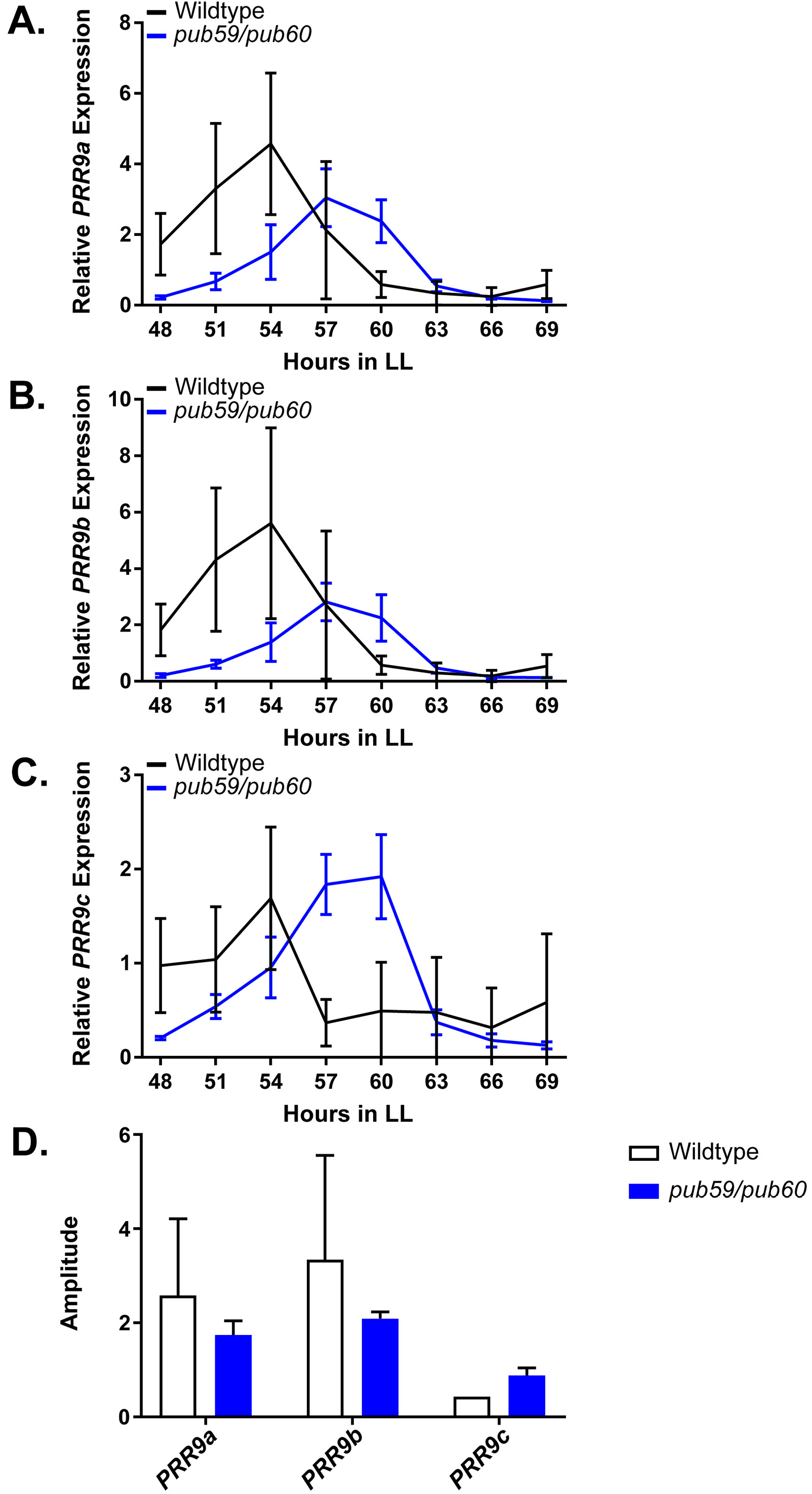
qRT-PCR of PRR9 splicing in *pub59/pub60* mutants. *PRR9* isoform expression was measured using quantitative RT-PCR *pub59/pub60* mutant lines. A) *PRR9a*, B) *PRR9b*, and C) *PRR9c* isoforms were analyzed. Quantifications are the average of three biological replicates with error bars showing standard deviation. D) FFT-NLLS analysis through the Biodare2 platform suggests altered isoform levels in the *pub59/pub60* double mutant. Error bars represent the standard deviation.

### Perturbations in *PUB60* expression lead to circadian clock defects

Accurate expression of genes involved in the circadian clock is essential to maintaining 24 hour periodicity (Más et al., 2003; Rawat et al., 2011; Somers et al., 2004). For this reason, we tested the effects of constitutive expression of full-length *PUB60* on circadian clock function. We created transgenic plants expressing FLAG-His-PUB60 under the control of a 35S constitutive promoter in the *CCA1p::Lucifernse* reporter line. Interestingly, constitutive expression of the full-length *PUB60* causes period lengthening similar to *pub59/pub60* double mutant (Fig 10). This indicates that maintaining proper expression of *PUB60* is necessary for clock function.

**Figure 10.**
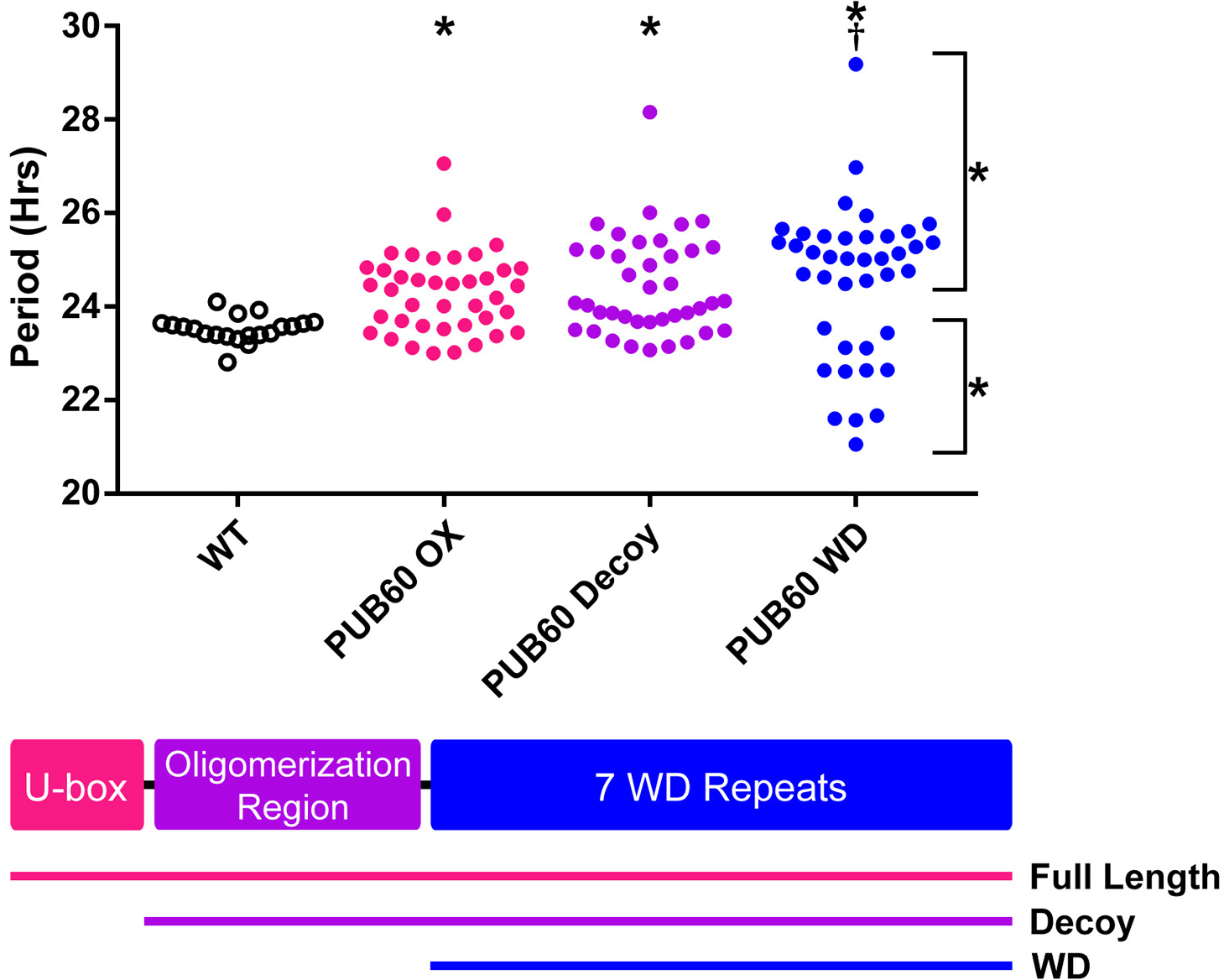
Period analyses of PUB60 overexpression constructs. Period was measured in T1 *PUB60* full length, *PUB60* decoy, and *PUB60* WD insertion lines. Period values presented are raw period values measured by *CCA1p::Luciferase* expression. A schematic of which domains are in each construct is included below.

### PUB60 decoys form biologically relevant complexes

We have previously shown that F-box decoy proteins are able to interact with target proteins and regulatory partners and retain the ability to form biologically relevant complexes (Lee and Feke et al., 2018). We tested whether the U-box decoy proteins are similarly capable of interacting with the same proteins as the full length U-box proteins.

The decoy proteins contain a 3XFLAG-6XHis affinity tag for immunoprecipitation, thus we performed an immunoprecipitation experiment with the PUB60 decoy and analyzed interacting proteins via mass spectrometry (IP-MS). As control we performed immunoprecipitation with a 3XFLAG-6XHis tagged GFP transgenic line. From the list of potential interacting proteins (Table S2) we identified known components of the NTC complex. We compared this list to the previously identified components of the plant NTC, and identified five common components in addition to the PUB60 bait peptides, three of which (PUB59, CELL DIVISION CYCLE 5 (CDC5), and MODIFIER OF SNC1,4 (MOS4)) were only found in the PUB60 IP-MS experiments and not in the controls (Table 4) (Monaghan et al., 2009). This data suggests that the PUB60 decoy is capable of forming biologically relevant complexes *in vivo* and importantly supports the idea that the decoy strategy can be used for genetic and biochemical analyses of E3 ligase function in plants.

**Table 4.**
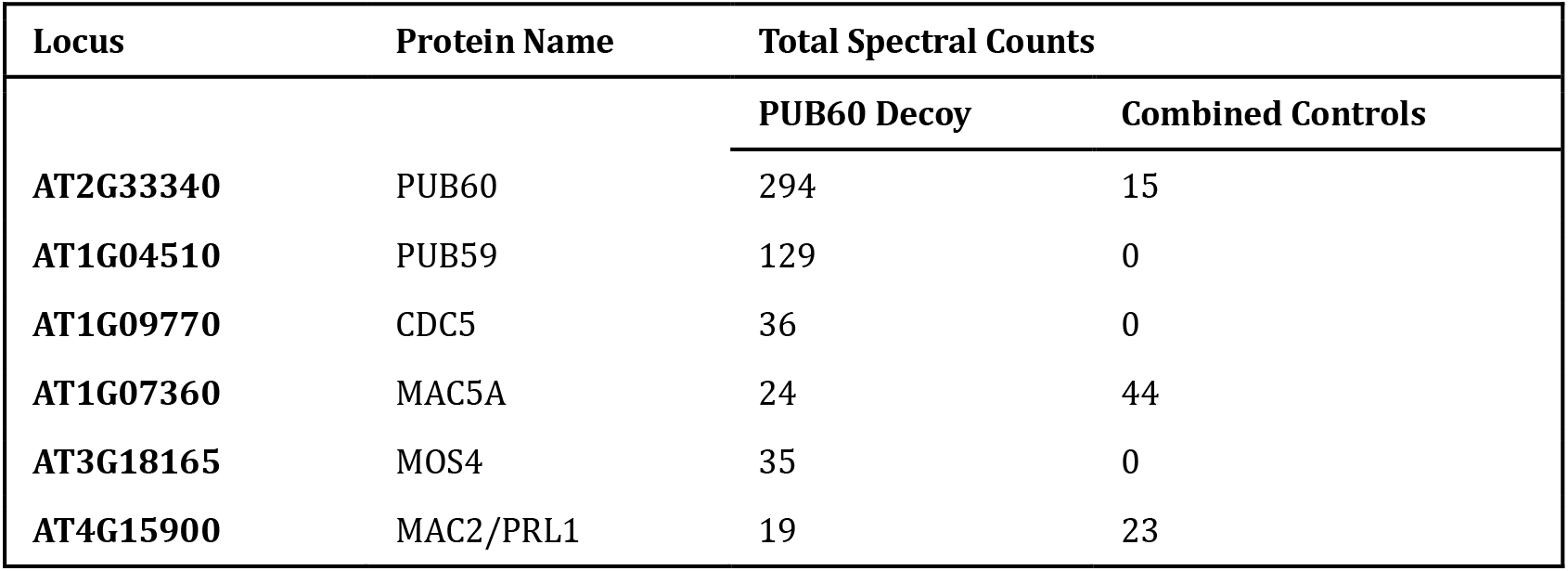
Selected IP-MS results from the PUB60 decoy. PUB60 decoy peptide hits are from one IP-MS experiment using the PUB60 decoy as the bait. Combined control peptide hits are summed from the independent control experiments of wildtype Col-0 and *35S::His-FLAG-GFP* expressing lines.

We observed PUB59 and PUB60 interacting in our IP/MS data and PRP19, the PUB59 and PUB60 orthologue from yeast and humans, is predicted to form a tetramer (Grillari et al., 2005; Ohi et al., 2005). Tetramer formation is predicted to be mediated by a conserved coiled coil that is present in PUB59 and PUB60 (Li et al., 2018), and was included in our PUB59 and PUB60 decoy constructs (Fig 10). We wanted to test the importance of the dimerization domain in clock function using our decoy system. Thus, we created a PUB60 decoy construct which expresses the WD repeats without the predicted oligomerization domain (PUB60 WD). Constitutive expression of the PUB60 WD domain causes two subpopulations of transgenic lines, one with lengthened period (25.5 hours) and one with shortened period (22.5 hours) (Fig 10). This was different than expressing the decoy or full-length PUB60 and indicates that oligomerization plays an important role in PUB60 function in the clock. Interestingly, the PUB60 WD phenotype more closely resembles the PUB59 decoy phenotype, suggesting that PUB59 and PUB60 may have diverged slightly in their biochemical function or oligomerization capabilities.

## DISCUSSION

### Summary

To overcome genetic redundancy we performed a large-scale reverse genetic screen of plant E3 ubiquitin ligases. We generated transgenic plants expressing dominant-negative E3 ubiquitin ligase decoys and determined the effects on the circadian clock. From this screen we identified the first U-box-type E3 ubiquitin ligases, PUB59 and PUB60, involved in circadian clock function along with additional putative “major” and “minor” clock period and phase regulators. Importantly, our detailed follow-up studies show that *PUB59* and *PUB60* are redundant regulators of clock function making it unlikely that they would be identified using traditional forward genetic screening methods. These genes represent the second family of E3 ubiquitin ligases to be identified in circadian clock function in plants, and their discovery uncovers a connection between three large cellular networks: the circadian clock, the ubiquitin proteasome, and splicing.

### PUB59 and PUB60 are part of the NTC and effect clock splicing

Splicing is a critical regulatory step in plant circadian clock function, and mutation of another component of the NTC, *SKIP*, lengthens circadian similar to the *pub59/pub60* double mutant. The lengthened period in the *skip* mutant is likely due to the dysregulation of *PRR9* and *PRR7* splicing (Wang et al., 2012). In concordance, we show that the *pub59/pub60* double mutant has defects in *PRR9* splicing, but it is also likely that other clock genes may have spliceoform imbalances in the *pub59/pub60* mutant as well (Jia et al., 2017).

*PUB59* and *PUB60* were identified as interacting partners of MOS4, a positive regulator of plant innate immunity. Genetic experiments then demonstrated that they are also required for plant immunity (Monaghan et al., 2009). They were named MOS4-Associated-Complex 3A and B (MAC3A and MAC3B) but share homology with the spliceosome activating component PRP19. In both yeast and humans, PRP19 is known to be the core component of a large complex, known as the NTC. The NTC plays key roles in DNA repair, cell cycle progression and genome maintenance, and activating the spliceosome (Chanarat and Sträßer, 2013; Hogg et al., 2010). Further work has since shown that *PUB59* and *PUB60* (alternatively *MAC3A* and *MAC3B* or *PRP19a* and *PRP19b*) are involved in the global regulation of splicing in plants (Jia et al., 2017). These results, along with the identification of plant NTC components interacting with PUB59 and PUB60 in this study and others, strongly suggest that these genes are core components of the plant NTC (Monaghan et al., 2009).

Because the clock is an interlocked series of feedback loops it can be difficult to predict how mutation or misexpression of a gene will affect clock function. Interestingly, this is the same for *PUB60* in which constitutive expression of *PUB60* results in a similar period defect as the *pub59/pub60* double mutant. There are two possible explanations: 1) PUB59 and PUB60 are involved in splicing various clock genes and the cumulative effect of their disruption is a lengthened period, or 2) splicing of clock genes is both positively and negatively regulated and disruption of this balance causes lengthened period. It is possible to distinguish between these two events by determining the full extent of clock gene mis-splicing in the *pub59/pub60* and *PUB60* overexpression lines, and then performing complementation experiments with the cDNA of the mis-spliced clock genes. Evidence from the *PUB59* decoy and *PUB60* WD populations indicates that it may be the former, as we observed short period lines in both populations. The absence of short period lines in the *PUB60* decoy is interesting, and suggests that these two proteins may have diverged slightly in their functions. Domain swap experiments or conversion of key residues between PUB59 and PUB60 may further elucidate these differences.

### The decoy technique uncovers difficult-to-identify clock regulators by overcoming redundancy

Our genetic studies of *PUB59* and *PUB60* highlight a critical strength of the decoy technique as a screening platform, the ability to overcome genetic redundancy. We observe minimal effect on the circadian clock in single *pub59* and *pub60* mutants. Yet, the double mutant has a strong non-additive genetic effect, suggesting the *PUB59* and *PUB60* genes are redundant. Exhaustive traditional forward genetic screens for clock mutants have not identified *PUB59* and *PUB60* (Hazen et al., 2005; Jones et al., 2012; Kevei et al., 2006; Martin-Tryon et al., 2006; Somers et al., 2000), suggesting that reverse genetic strategies, such as our decoy approach, are important for generating a comprehensive genetic picture of the plant circadian clock.

### Description of candidate clock genes

In addition to *PUB59* and *PUB60*, we also identified 43 F-box and U-box genes which are likely to be involved in the regulation of the phase circadian clock, and 42 genes which are likely to be involved in the regulation of the period. We have highlighted seven of these as high priority regulators due to their strong phenotypic effects. These genes include *ACF1 (AT5G44980), ACF2 (AT5G48980), CFK1, ACF3 (AT1G20800), ACF4 (AT2G44030), ACF5* (.*AT1G50870*), *PUB48*, and *PUB30*.

Nearly nothing is known about four of the high-priority *ACFs* that we discovered in our study. While future work will be required to untangle the relationships between these three genes and the circadian clock, our ability to isolate these novel genes highlights the strengths of the decoy library. All three genes have close homologs (Table 2), so it is possible that, similar to *PUB59* and *PUB60*, forward genetic screens failed to identify these genes due to redundancy.

*CFK1* is a regulator of hypocotyl length, and *CFK1* expression is light-induced (Franciosini et al., 2013). Light is known to control both the phasing of the circadian clock as well as hypocotyl elongation, so the ability of *CFK1* to respond to light signals makes it a promising candidate. Furthermore, *CFK1* is believed to be a target of the COP9 signalosome (CSN). The CSN is required for proper rhythmicity in *Neurospora*, and CSN mutants lead to impaired phase resetting in *Drosophila* (He et al., 2005; Knowles et al., 2009; Zhou et al., 2012). It is possible that the CSN plays a similar role in plants, possibly through regulation of *CFK1*. Future work detailing these connections between *CFK1*, the CSN, and the plant circadian clock would likely be quite fruitful.

In our U-box screen functions for putative hits are partially understood. *PUB48* and *PUB30* are involved in stress responses. *PUB30* is involved in the response to high salt conditions, and mutants exhibit reduced salt tolerance (Hwang et al., 2015; Zhang et al., 2017). Perturbations in the circadian clock can lead to altered salt stress response, so it is possible that alterations in the circadian clock of the *pub30* mutants similar to what we observe with the *PUB30* decoy leads to the observed salt stress phenotype (Kim et al., 2013; Nakamichi et al., 2009). *PUB48* is involved in response to drought stress (Adler et al., 2017). Although drought tolerance has not been implicated as an input to the circadian clock, water intake has been demonstrated to be a clock output (Takase et al., 2011). It is possible that altering the circadian clock in these plants may lead to improper regulation of water intake, and thus cause the drought stress sensitivity phenotype. Alternatively, recent work demonstrates that humidity cycles are sufficient to entrain the plant circadian clock, and contributes to rhythmicity even under cycling light conditions (Mwimba et al., 2018).

*PUB48* may play a role in regulating this input pathway, and thus lead to both altered drought sensitivity and an altered circadian clock. Future work on investigating the intersection between PUB48, water levels, and the circadian clock should help illuminate these connections.

All of the candidate regulators were identified using our reverse genetic decoy approach. It is imperative that further genetic and molecular work be performed to confirm their roles in clock function. Full-length overexpression or mutant studies would be informative; however, as demonstrated by our work on *PUB59* and *PUB60*, higher order mutants may be needed in order to uncover the functions of redundant genes. It is our hope that the candidate genes identified in this manuscript will serve as a springboard for future work in the field.

### Conclusions

The data presented in this manuscript demonstrates that decoys are a potent and scalable technique for identifying the function of plant E3 ubiquitin. The molecular and genetic reagents generated in the course of this study are already available to the community. The library that we have created will be available in multiple formats, including vectors (Gateway compatible entry vectors, the HIS-FLAG tagged expression vectors), agrobacterium for stable transformation into a mutant line of interest or transient expression, and a transgenic seed collection available in *pCCA1::Lucifernse* or Col-0). Furthermore, we have shown here that this technique applies to multiple families of E3 ubiquitin ligases, the U-boxes and the F-boxes. It is a logical extension to believe this technique would work for other families of E3 ubiquitin ligases, so long as the domains involved in interaction with the E2 conjugating enzyme are easily defined. We believe that the data shown here demonstrates that the decoy technique is a valuable resource to anyone interested in uncovering the function of plant E3 ubiquitin ligases involved in any aspect of plant biology.

## METHODS

### Construction of Decoy Libraries

In order to create F-box and U-box decoys, the CDS annotation from TAIR10 was compared to the protein domain annotation from Uniprot, and the nucleotide boundaries for the start and end of the E3 ubiquitin ligase domain were recorded. As the majority of F-box proteins follow a standardized domain architecture with the F-box domain located in the N-terminus of the protein, decoy constructs were created by removing the nucleotides of the F-box and any nucleotides upstream of the F-box domain. The U-box proteins, however, do not follow a standardized domain architecture, as the U-box domain can be located anywhere throughout the protein. For this reason, for the purposes of cloning the decoy library, U-boxes were sorted into three categories: N-terminal U-boxes, C-terminal U-boxes, and central U-boxes. Those genes where the U-box domain began less than 75 amino acids from the start of the protein were sorted into the N-terminal U-box class; those genes where the U-box domain ends less than 75 amino acids from the end of the protein were sorted into the C-terminal U-box class; and those genes where the U-box domain begins more than 75 amino acids from the beginning of the protein and ends more than 75 amino acids from the end of the protein were sorted into the central U-box class. 75 amino acids was selected as the threshold as there were gaps in U-box distribution throughout the protein sequence which made this a natural choice.

Primers for creation of F-box and U-box decoys were designed using the CDS annotation from TAIR10 (Table S3). For central U-boxes, N-terminal and C-terminal constructs were generated using PCR products generated from cDNA, then overlap extension PCR was used to fuse N-terminal and C-terminal constructs into the full decoy construct. PCR products generated from cDNA were inserted into pENTR/D-TOPO vectors (Invitrogen, cat. # K240020) then transferred into pB7-HFN, pK7-HFN, and pB7-HFC destination vectors using LR recombination (Huang et al., 2016a). F-boxes and N-terminal U-boxes were cloned into pB7-HFN and pK7-HFN (N-terminal His-FLAG tags), C-terminal U-boxes were cloned into pB7-HFC (C-terminal HIS-FLAG tags), and central U-boxes were cloned into all three vectors. The decoy constructs were transformed into Arabidopsis Col-0 expressing the circadian reporter *pCCA1::Lucifernse* or Col-0 by the floral dip method using *Agrobacterium tumefaciens* GV3101 (Pruneda-Paz et al., 2009).

### Phenotypic Screening

Control *pCCA1::Lucifernse* and decoy seeds were surface sterilized in 70% ethanol and 0.01% Triton X-100 for 20 minutes prior to being sown on ½ MS plates (2.15 g/L Murashige and Skoog medium, pH 5.7, Cassion Laboratories, cat#MSP01 and 0.8% bacteriological agar, AmericanBio cat# AB01185) with or without appropriate antibiotics (15 μg/ml ammonium glufosinate (Santa Cruz Biotechnology, cat# 77182-82-2) for vectors pB7-HFN and pB7-HFC, or 50 μg/ml kanamycin sulfate (AmericanBio) for pK7-HFN). Seeds were stratified for two days at 4 °C, then transferred to 12 hr light/12 hr dark conditions for seven days. Seven-day old seedlings were arrayed on 100 mm square ½ MS plates in a 10×10 grid, then treated with 5 mM D-luciferin (Cayman Chemical Company, cat# 115144-35-9) dissolved in 0.01% TritonX-100. Seedlings were imaged at 22 °C under constant white light provided by two LED light panels (Heliospectra L1) with light fluence rate of 100 μmol m^-2^ s^-1^. The imaging regime is as follows: each hour lights are turned off for two minutes, then an image is collected using a five minute exposure on an Andor iKon-M CCD camera; lights remain off for one minute after the exposure is completed, then lights return to the normal lighting regime. The CCD camera was controlled using Micromanager, using the following settings: binning of 2, pre-am gain of 2, and a 0.05 MHz readout mode (Edelstein et al., 2014). Using this setup, 400 seedlings are simultaneously imaged across four plates. Images are acquired each hour for approximately six and a half days. Data collected between the first dawn of constant light and the dawn of the sixth day are used for analyses.

The mean intensity of each seedling at each time point was calculated using ImageJ (Schneider et al., 2012). The calculated values were imported into the Biological Rhythms Analysis Software System (BRASS) for analysis. The Fast Fourier Transform Non-linear Least Squares (FFT-NLLS) algorithm was used to calculate the period, phase, and relative amplitude from each individual seedling (Moore et al., 2014).

### Data Normalization and Statistical Analysis

To allow for comparison across independent imaging experiments, period and phase data was normalized to the individual wildtype control performed concurrently. The average value of the wildtype control lines was calculated for every experiment, then this average was subtracted from the value of each individual T1 insertion or control wildtype line done concurrently. This normalized value was used for statistical analyses.

The presence of sub-populations was determined by a custom MATLAB script which takes the normalized values as inputs and creates histograms of each population. The peaks and troughs of the histogram are identified, and number of seedlings within each peak (between each pair of troughs) was counted. Peaks were discarded if the number of seedlings was too small (less than 3), if there was only one bin between peaks, or if the difference between peak and trough was too small (less than 3). If the number of remaining peaks was two or more, the population was defined as having subpopulations. The locations of the troughs in the histogram were used as the division point to sort lines into their respective subpopulations.

Welch’s t-test was used to compare each normalized T1 insertion line population or subpopulation to the population of normalized control lines. For period, all wildtype lines were used as the control. For phase, the entire population of decoy lines was used as the control. Data from the F-box decoy library was treated as an independent experiment from data from the U-box decoy library. In order to decrease the number of false positives caused by multiple testing, we utilized a Bonferroni corrected α as the p-value threshold. The α applied differs between experiments, and is noted throughout.

### Measurement of Circadian Gene Expression in *pub59/pub60* mutant lines

Homozygous *pub59/pub60* mutant lines in the Col-0 background were generated previously (Monaghan et al., 2009). Single and double *pub59* and *pub60* mutant and Col-0 seeds were grown on ½ MS plates and entrained in 12 hr light/12 hr dark conditions at a fluence rate of 130 μmol m^-2^ s^-1^ at 22 °C. 10-day old seedlings were transferred into constant light conditions for 48 hours prior to the start of the time course. Seedlings were collected every three hours for two days starting at ZT0 and snap-frozen using liquid nitrogen, then ground using the Mixer Mill MM400 system (Retsch). Total RNA was extracted from ground seedlings using the RNeasy Plant Mini Kit and treated with RNase-Free DNase (Qiagen, cat#74904 and 79254) following the manufacturer’s protocols. cDNA was prepared from 1 μg total RNA using iScript™ Reverse Transcription Supermix (BioRad, cat#1708841), then diluted 15-fold and used directly as the template for quantitative real-time RT-PCR (qRT-PCR). The qRT-PCR was performed using 3.5 μl of diluted cDNA and 5.5 μM primers listed in Table S3 (Czechowski et al., 2004; Farré et al., 2005; Lee and Thomashow, 2012)using iTaq™ Universal SYBR^®^ Green Supermix (Bio-Rad, cat# 1725121) with the CFX 384 Touch™ Real-Time PCR Detection System (Bio-RAD). The qRT-PCR began with a denaturation step of 95°C for 3 min, followed by 45 cycles of denaturation at 95°C for 15 sec, and primer annealing at 53°C for 15s. Relative expression of *CCA1* and *TOC1* was determined by the comparative C_T_ method using *IPP2 (AT3G02780*) as an internal control. The relative expression levels represent the mean values of 2^-ΔΔCT^ from three biological replicates, where ΔCT = C_T_ of the decoy – C_T_ IPP2 and the reference point is the first peak time for each replicate (ZT0 for Col-0, *pub59*, and *pub60*, and ZT6 for *pub59/pub60* for *CCA1* expression, and ZT12 for Col-0, *pub59*, and *pub60*, and ZT15 for *pub59/60* for *TOC1* expression).

### Measurement of PRR9 spliceoforms

*pub59/pub60* double mutant seedlings in the Col-0 background and parental Col-0 seeds were grown and harvested as described for circadian gene expression analysis. qPCR was performed as described for *CCA1* expression analysis. Primers used in a previous study (Wang et al., 2012) to track *PRR9* spliceoform expression are shown in Table S3. The relative expression levels represent the mean values of 2^-ΔΔCT^ from three biological replicates, where ΔCT = C_T_ of the decoy – C_T_ IPP2 and the reference point is ZT0 from one of the Col-0 replicates.

### Immunoprecipitation and Mass Spectrometry of PUB60 Decoy lines

Individual T1 *pB7-HFN-PUB60* transgenic lines in a Col-0 background and control Col-0 and *pB7-HFC-GFP* were grown as described for phenotype analysis. Seven-day old seedlings were transferred to soil and grown under 16 hours light/8 hours dark at 22 °C for 2-3 weeks. Prior to harvest, plants were entrained to 12 hours light/12 hours dark at 22 °C for 1 week. Approximately 40 mature leaves from each background was collected and flash frozen in liquid nitrogen, such that each sample was a mixture of leaves from multiple individuals to reduce the effects of expression level fluctuations. Tissue samples were ground in liquid nitrogen using the Mixer Mill MM400 system (Retsch). Immunoprecipitation was performed as described previously (Huang et al., 2016a, 2016b; Lu et al., 2010). Briefly, protein from 2 ml tissue powder was extracted in SII buffer (100mM sodium phosphate pH 8.0, 150 mM NaCl, 5 mM EDTA, 0.1% Triton X-100) with cOmplete™ EDTA-free Protease Inhibitor Cocktail (Roche, cat# 11873580001), 1 mM phenylmethylsμlfonyl fluoride (PMSF), and PhosSTOP tablet (Roche, cat# 04906845001) by sonification. Anti-FLAG antibodies were cross-linked to Dynabeads^®^ M-270 Epoxy (Thermo Fisher Scientific, cat# 14311D) for immunoprecipitation. Immunoprecipitation was performed by incubation of protein extracts with beads for 1 hour at 4 °C on a rocker. Beads were washed with SII buffer three times, then twice in F2H buffer (100 mM sodium phosphate pH 8.0, 150 mM NaCl, 0.1% Triton X-100). Beads were eluted twice at 4 °C and twice at 30 °C in F2H buffer with 100 μg/ml FLAG peptide, then incubated with TALON magnetic beads (Clontech, cat# 35636) for 20 min at 4 °C, then washed twice in F2H buffer and three times in 25 mM Ammonium Bicarbonate. Samples were subjected to trypsin digestion (0.5 μg, Promega, cat# V5113) at 37 °C overnight, then vacuum dried using a SpeedVac before being dissolved in 5% formic acid/0.1% trifluoroacetic acid (TFA). Protein concentration was determined by nanodrop measurement (A260/A280)(Thermo Scientific Nanodrop 2000 UV-Vis Spectrophotometer). An aliquot of each sample was further diluted with 0.1% TFA to 0.1μg/μl and 0.5μg was injected for LC-MS/MS analysis at the Keck MS & Proteomics Resource Laboratory at Yale University.

LC-MS/MS analysis was performed on a Thermo Scientific Orbitrap Elite mass spectrometer equipped with a Waters nanoAcquity UPLC system utilizing a binary solvent system (Buffer A: 0.1% formic acid; Buffer B: 0.1% formic acid in acetonitrile). Trapping was performed at 5μl/min, 97% Buffer A for 3 min using a Waters Symmetry^®^ C18 180μm x 20mm trap column. Peptides were separated using an ACQUITY UPLC PST (BEH) C18 nanoACQUITY Column 1.7 μm, 75 μm x 250 mm (37°C) and eluted at 300 nl/min with the following gradient: 3% buffer B at initial conditions; 5% B at 3 minutes; 35% B at 140 minutes; 50% B at 155 minutes; 85% B at 160-165 min; then returned to initial conditions at 166 minutes. MS were acquired in the Orbitrap in profile mode over the 300-1,700 m/z range using 1 microscan, 30,000 resolution, AGC target of 1E6, and a full max ion time of 50 ms. Up to 15 MS/MS were collected per MS scan using collision induced dissociation (CID) on species with an intensity threshold of 5,000 and charge states 2 and above. Data dependent MS/MS were acquired in centroid mode in the ion trap using 1 microscan, AGC target of 2E4, full max IT of 100 ms, 2.0 m/z isolation window, and normalized collision energy of 35. Dynamic exclusion was enabled with a repeat count of 1, repeat duration of 30s, exclusion list size of 500, and exclusion duration of 60s.

The MS/MS spectra were searched by the Keck MS & Proteomics Resource Laboratory at Yale University using MASCOT (Perkins et al., 1999). Data was searched against the SwissProt_2015_11.fasta *Arabidopsis thaliana* database with oxidation set as a variable modification. The peptide mass tolerance was set to 10 ppm, the fragment mass tolerance to 0.5 Da, and the maximum number of allowable missed cleavages was set to 2.

**Figure S1.**
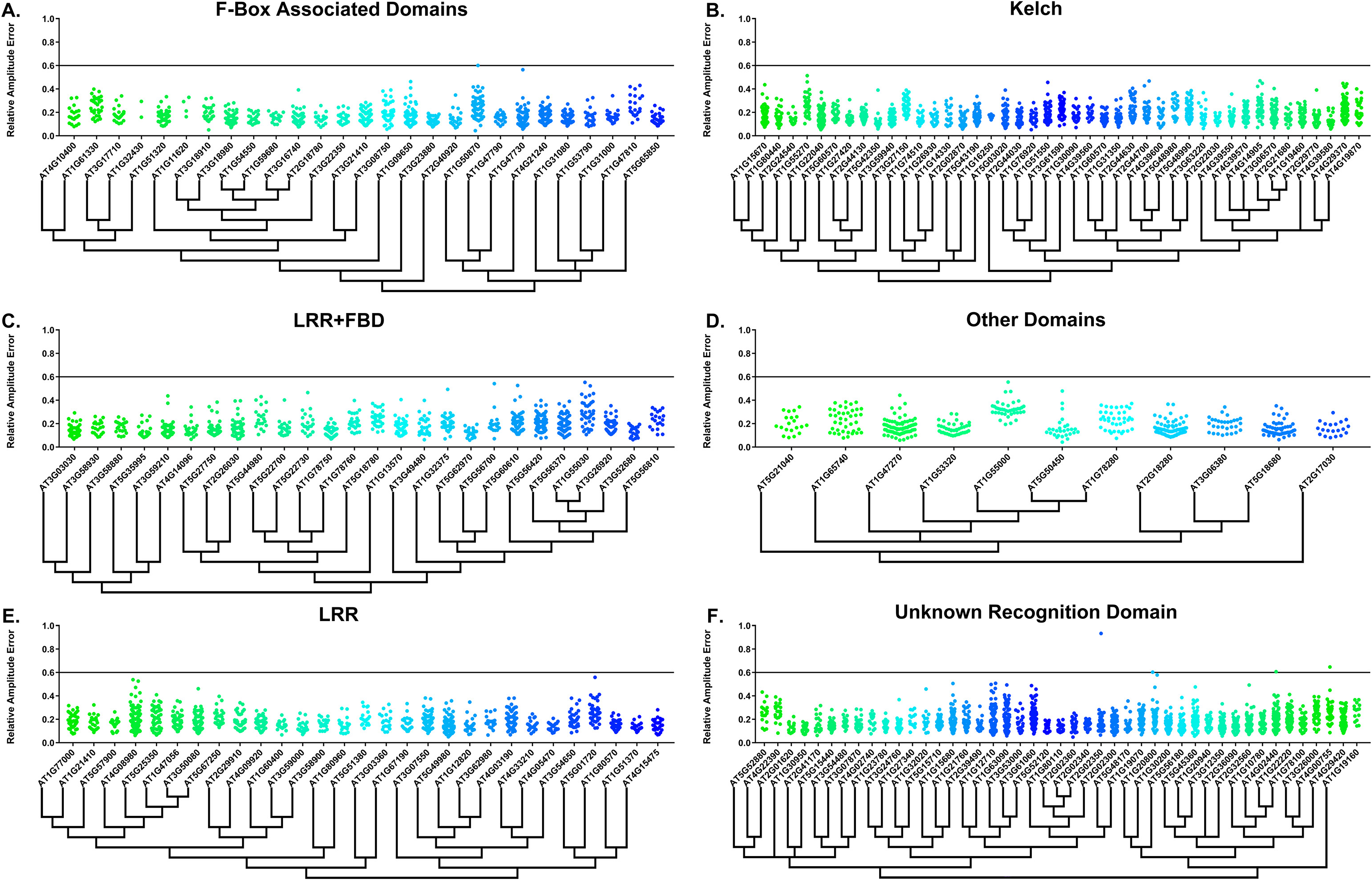
RAE distributions of F-box decoy lines. Values presented are the RAE for each individual T1 seedling. The black line represents the standard RAE cutoff of 0.6. Genes are separated by protein recognition domain and ordered by closest protein homology using Phylogeny.Fr, (Dereeper et al., 2008), and a tree showing that homology is displayed beneath the graph. F-Box Associated Domains = FBA1, FBA3, and FBD only. Other Domains = TUB, JmjC, LysM, WD40, zf_MYND, and DUF295. Only data from those experiments where the control *CCA1p::Luciferaselines* display a standard deviation less than 0.75 were included in our analyses.

**Figure S2.**
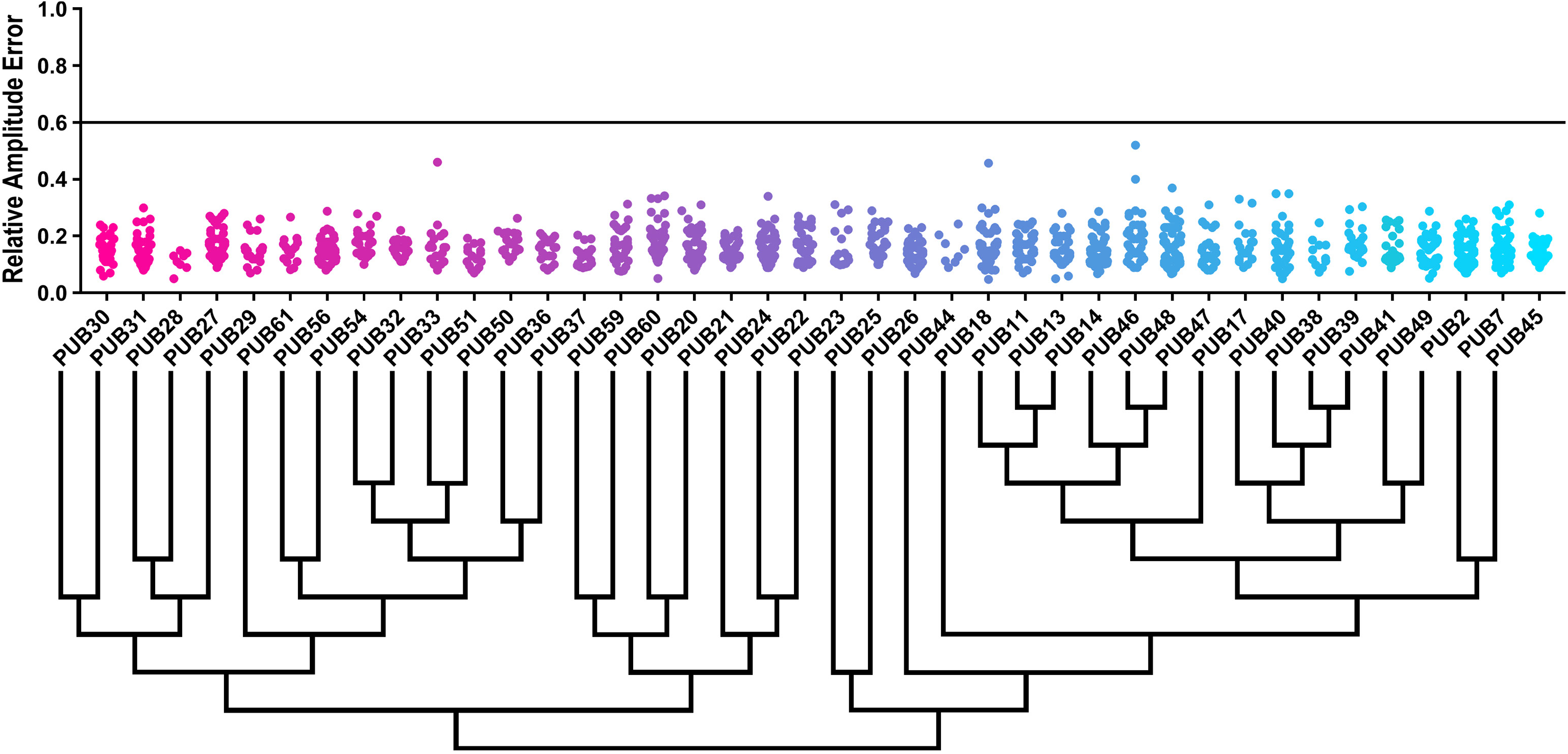
RAE distributions of U-box decoy lines. Values presented are the RAE for each individual T1 seedling. The black line represents the standard RAE cutoff of 0.6. Genes are ordered by closest protein homology using Phylogeny.Fr, (Dereeper et al., 2008), and a tree showing that homology is displayed beneath the graph.

**Figure S3.**
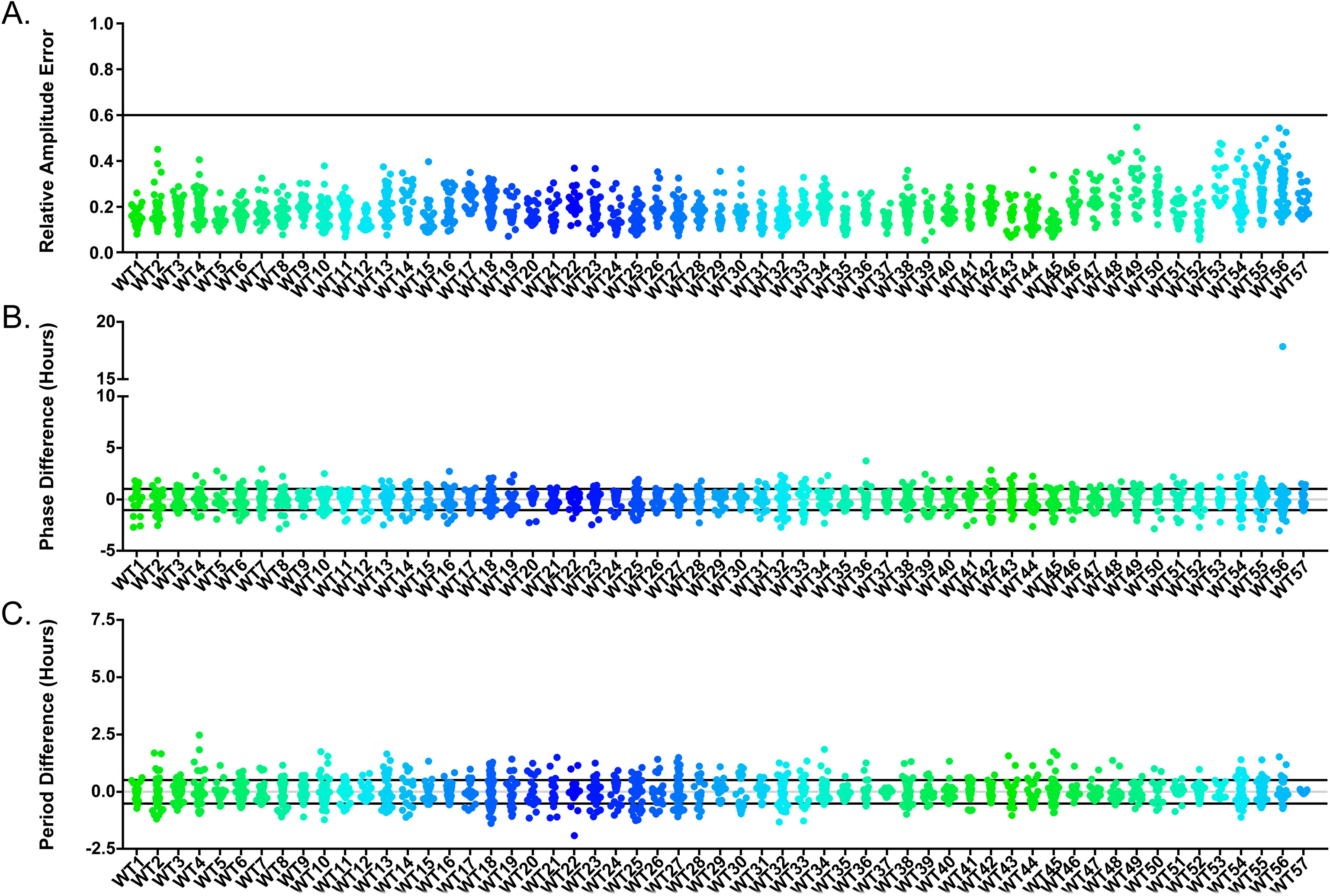
Circadian distributions of control lines in F-box experiments. Values presented are A) the RAE of each individual control line. The black line represents the standard RAE cutoff of 0.6. B) the difference between the phase of the individual control line and the average phase of the *CCA1p::Luciferase* control in the accompanying experiment. The grey line is at the average control value and the black lines are at +/– the standard deviation of the control lines. The grey line is at the average control value and the black lines are at +/– the standard deviation of the control lines. C) the difference between the period of the individual control line and the average period of the *CCA1p::Luciferase* control in the accompanying experiment. The grey line is at the average control value and the black lines are at +/– the standard deviation of the control lines. Only data from those experiments where the control *CCA1p::Luciferase* lines display a standard deviation less than 0.75 were included in our analyses.

**Figure S4.**
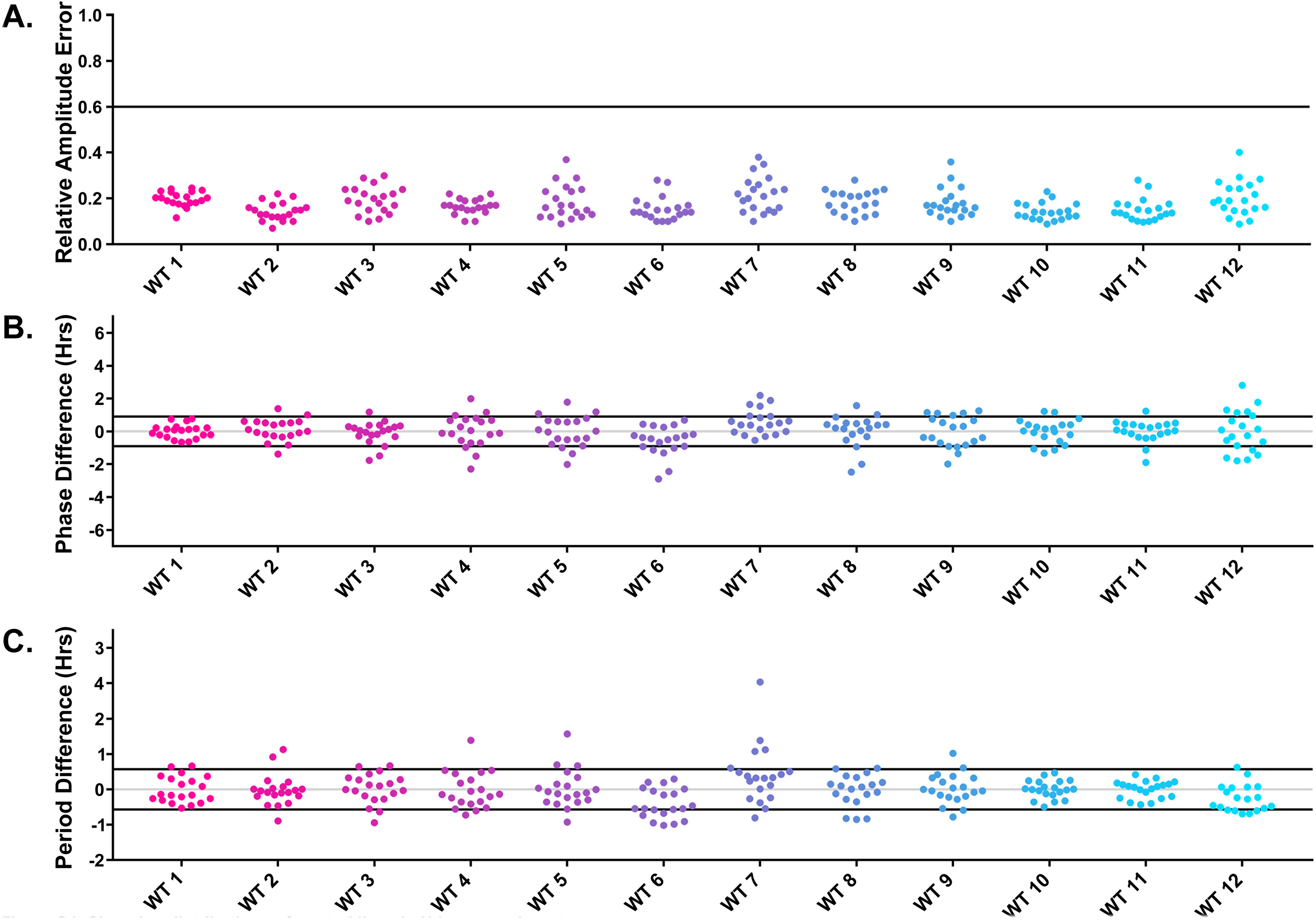
Circadian distributions of control lines in U-box experiments. Values presented are A) the RAE of each individual control line. The black line represents the standard RAE cutoff of 0.6. B) the difference between the phase of the individual control line and the average phase of the *CCA1p::Luciferase* control in the accompanying experiment. The grey line is at the average control value and the black lines are at +/– the standard deviation of the control lines. The grey line is at the average control value and the black lines are at +/– the standard deviation of the control lines. C) the difference between the period of the individual control line and the average period of the *CCA1p::Luciferase* control in the accompanying experiment. The grey line is at the average control value and the black lines are at +/– the standard deviation of the control lines.

Table S1. All generated data and publications which reference genes in our decoy library.

Table S2. IP-MS results from the PUB60 decoy.

Table S3. Primers used in this Study.

## Acknowledgements

We would like to thank Christopher Adamchek, Cathy Chamberlin, Suyuna Eng Ren, Sandra Pariseau, Denise George, Brandon Williams, Milan Sandhu, and Annie Jin for their technical support. We would also like to thank the Keck Proteomics Facility at Yale for their assistance with proteomics, and Dr. Xin Li for providing the *pub59* and *pub59/pub60* double mutants. Additionally, we would like to thank Dr. Qingqing Wang and Dr. Bryan Thines for their helpful comments on the manuscript.

## Contributions

JMG, AF, WL, CML, and MWL conceived of and designed the decoy libraries used in this study. AF, WL, JH, CML, MWL, and EZ cloned and created the decoy library. AF and JMG conceived of the experiments for PUB59 and PUB60. AF performed the experiments and experimental analyses. AF and JMG wrote the manuscript.

## REFERENCES

Adler G, Konrad Z, Zamir L, Mishra AK, Raveh D, Bar-Zvi D. 2017. The Arabidopsis paralogs, PUB46 and PUB48, encoding U-box E3 ubiquitin ligases, are essential for plant response to drought stress. BMC Plant Biol 17:8. doi:10.1186/s12870-016-0963-5

Alabadi D, Oyama T, Yanovsky MJ, Harmon FG, Mas P, Kay SA. 2001. Reciprocal Regulation Between TOC1 and LHY/CCA1 Within the Arabidopsis Circadian Clock. Science (80–) 293:880–883. doi:10.1126/science.1061320

Alabadí D, Yanovsky MJ, Mas P, Harmer SL, Kay SA. 2002. Critical role for CCA1 and LHY in maintaining circadian rhythmicity in Arabidopsis. Curr Biol 12:757–61.

Andersen P, Kragelund BB, Olsen AN, Larsen FH, Chua N-H, Poulsen FM, Skriver K. 2004. Structure and biochemical function of a prototypical Arabidopsis U-box domain. J Biol Chem 279:40053–61. doi:10.1074/jbc.M405057200

Andrade MA, González-Guzmán M, Serrano R, Rodríguez PL. 2001. A combination of the F-box motif and kelch repeats defines a large Arabidopsis family of F-box proteins. Plant Mol Biol 46:603–14.

Aravind L, Koonin E V. 2000. The U box is a modified RING finger – a common domain in ubiquitination. Curr Biol 10:R132–R134. doi:10.1016/S0960-9822(00)00398-5

Azevedo C, Santos-Rosa MJ, Shirasu K. 2001. The U-box protein family in plants. Trends PlantSci 6:354–8.

Bai C, Sen P, Hofmann K, Ma L, Goebl M, Harper JW, Elledge SJ. 1996. SKP1 connects cell cycle regulators to the ubiquitin proteolysis machinery through a novel motif, the F-box. Cell 86:263–74.

Carré IA, Kim J-Y. 2002. MYB transcription factors in the Arabidopsis circadian clock. J Exp Bot 53:1551–7.

Chanarat S, Sträßer K. 2013. Splicing and beyond: The many faces of the Prp19 complex. Biochim Biophys Acta – Mol Cell Res 1833:2126–2134. doi:10.1016/j.bbamcr.2013.05.023

Chen L, Hellmann H. 2013. Plant E3 Ligases: Flexible Enzymes in a Sessile World. Mol Plant 6:1388–1404. doi:10.1093/mp/sst005

Czechowski T, Bari RP, Stitt M, Scheible W-R, Udvardi MK. 2004. Real-time RT-PCR profiling of over 1400 *Arabidopsis* transcription factors: unprecedented sensitivity reveals novel root– and shoot-specific genes. Plant J 38:366–379. doi:10.1111/j.1365-313X.2004.02051.x

Dereeper A, Guignon V, Blanc G, Audic S, Buffet S, Chevenet F, Dufayard J-F, Guindon S, Lefort V, Lescot M, Claverie J-M, Gascuel O. 2008. Phylogeny.fr: robust phylogenetic analysis for the non-specialist. Nucleic Acids Res 36:W465–W469. doi:10.1093/nar/gkn180

Deshaies RJ. 1999. SCF and Cullin/Ring H2-based ubiquitin ligases. Annu Rev Cell Dev Biol 15:435–67. doi:10.1146/annurev.cellbio.15.1.435

Deshaies RJ, Joazeiro CAP. 2009. RING domain E3 ubiquitin ligases. Annu Rev Biochem 78:399–434. doi:10.1146/annurev.biochem.78.101807.093809

Dowson-Day MJ, Millar AJ. 1999. Circadian dysfunction causes aberrant hypocotyl elongation patterns in Arabidopsis. Plant J 17:63–71.

Doyle MR, Davis SJ, Bastow RM, McWatters HG, Kozma-Bognar L, Nagy F, Millar AJ, Amasino RM. 2002. The ELF4 gene controls circadian rhythms and flowering time in Arabidopsis thaliana. Nature 419:74–77. doi:10.1038/nature00954

Edelstein AD, Tsuchida MA, Amodaj N, Pinkard H, Vale RD, Stuurman N. 2014. Advanced methods of microscope control using μManager software. J Biol methods 1:10. doi:10.14440/jbm.2014.36

Endo M, Shimizu H, Nohales MA, Araki T, Kay SA. 2014. Tissue-specific clocks in Arabidopsis show asymmetric coupling. Nature 515:419–422. doi:10.1038/nature13919

Farinas B, Mas P. 2011. Histone acetylation and the circadian clock: a role for the MYB transcription factor RVE8/LCL5. Plant Signal Behav 6:541–3.

Farré EM, Harmer SL, Harmon FG, Yanovsky MJ, Kay SA. 2005. Overlapping and Distinct Roles of PRR7 and PRR9 in the Arabidopsis Circadian Clock. Curr Biol 15:47–54. doi:10.1016/j.cub.2004.12.067

Filichkin SA, Mockler TC. 2012. Unproductive alternative splicing and nonsense mRNAs: A widespread phenomenon among plant circadian clock genes. Biol Direct 7:20. doi:10.1186/1745-6150-7-20

Finn RD, Coggill P, Eberhardt RY, Eddy SR, Mistry J, Mitchell AL, Potter SC, Punta M, Qureshi M, Sangrador-Vegas A, Salazar GA, Tate J, Bateman A. 2016. The Pfam protein families database: towards a more sustainable future. Nucleic Acids Res 44:D279–D285. doi:10.1093/nar/gkv1344

Fowler S, Lee K, Onouchi H, Samach A, Richardson K, Morris B, Coupland G, Putterill J. 1999. GIGANTEA: a circadian clock-controlled gene that regulates photoperiodic flowering in Arabidopsis and encodes a protein with several possible membrane-spanning domains. EMBO J 18:4679–88. doi:10.1093/emboj/18.17.4679

Franciosini A, Lombardi B, Iafrate S, Pecce V, Mele G, Lupacchini L, Rinaldi G, Kondou Y, Gusmaroli G, Aki S, Tsuge T, Deng X-W, Matsui M, Vittorioso P, Costantino P, Serino G. 2013. The Arabidopsis COP9 SIGNALOSOME INTERACTING F-BOX KELCH 1 Protein Forms an SCF Ubiquitin Ligase and Regulates Hypocotyl Elongation. Mol Plant 6:1616–1629. doi:10.1093/mp/sst045

Fujimore T, Sato E, Yamashino T, Mizuno T. 2005. PRR5 (PSEUDO-RESPONSE REGULATOR 5) Plays Antagonistic Roles to CCA1 (CIRCADIAN CLOCK-ASSOCIATED 1) in Arabidopsis thaliana. Biosci Biotechnol Biochem 69:426–430. doi:10.1271/bbb.69.426

Fujiwara S, Wang L, Han L, Suh S-S, Salome PA, McClung CR, Somers DE. 2008. Post-translational regulation of the Arabidopsis circadian clock through selective proteolysis and phosphorylation of pseudo-response regulator proteins. J Biol Chem 283:23073–83. doi:10.1074/jbc.M803471200

Gendron JM, Pruneda-Paz JL, Doherty CJ, Gross AM, Kang SE, Kay SA. 2012. Arabidopsis circadian clock protein, TOC1, is a DNA-binding transcription factor. Proc Natl Acad Sci 109:3167–3172. doi:10.1073/pnas.1200355109

Godinho SIH, Maywood ES, Shaw L, Tucci V, Barnard AR, Busino L, Pagano M, Kendall R, Quwailid MM, Romero MR, O’neill J, Chesham JE, Brooker D, Lalanne Z, Hastings MH, Nolan PM. 2007. The after-hours mutant reveals a role for Fbxl3 in determining mammalian circadian period. Science 316:897–900. doi:10.1126/science.1141138

Grillari J, Ajuh P, Stadler G, Löscher M, Voglauer R, Ernst W, Chusainow J, Eisenhaber F, Pokar M, Fortschegger K, Grey M, Lamond AI, Katinger H. 2005. SNEV is an evolutionarily conserved splicing factor whose oligomerization is necessary for spliceosome assembly. Nucleic Acids Res 33:6868–6883. doi:10.1093/nar/gki986

Grima B, Lamouroux A, Chélot E, Papin C, Limbourg-Bouchon B, Rouyer F. 2002. The F-box protein slimb controls the levels of clock proteins period and timeless. Nature 420:178–82. doi:10.1038/nature01122

Han L, Mason M, Risseeuw EP, Crosby WL, Somers DE. 2004. Formation of an SCF(ZTL) complex is required for proper regulation of circadian timing. Plant J 40:291–301. doi:10.1111/j.1365-313X.2004.02207.x

Hazen SP, Schultz TF, Pruneda-Paz JL, Borevitz JO, Ecker JR, Kay SA. 2005. LUX ARRHYTHMO encodes a Myb domain protein essential for circadian rhythms. Proc Natl Acad Sci 102:10387–10392. doi:10.1073/pnas.0503029102

He Q, Cheng P, He Q, Liu Y. 2005. The COP9 signalosome regulates the Neurospora circadian clock by controlling the stability of the SCFFWD-1 complex. Genes Dev 19:1518–31. doi:10.1101/gad.1322205

He Q, Cheng P, Yang Y, He Q, Yu H, Liu Y. 2003. FWD1-mediated degradation of FREQUENCY in Neurospora establishes a conserved mechanism for circadian clock regulation. EMBO J 22:4421–30. doi:10.1093/emboj/cdg425

Helfer A, Nusinow DA, Chow BY, Gehrke AR, Bulyk ML, Kay SA. 2011. LUX ARRHYTHMO Encodes a Nighttime Repressor of Circadian Gene Expression in the Arabidopsis Core Clock. Curr Biol 21:126–133. doi:10.1016/j.cub.2010.12.021

Hicks KA, Albertson TM, Wagner DR. 2001. EARLY FLOWERING3 encodes a novel protein that regulates circadian clock function and flowering in Arabidopsis. Plant Cell 13:1281–92.

Hicks KA, Millar AJ, Carré IA, Somers DE, Straume M, Meeks-Wagner DR, Kay SA. 1996. Conditional circadian dysfunction of the Arabidopsis early-flowering 3 mutant. Science 274:790–2.

Hogg R, McGrail JC, O’Keefe RT. 2010. The function of the NineTeen Complex (NTC) in regulating spliceosome conformations and fidelity during pre-mRNA splicing. Biochem Soc Trans 38:1110–1115. doi:10.1042/BST0381110

Hoshizaki T, Hamner KC. 1964. Circadian Leaf Movements: Persistence in Bean Plants Grown in Continuous High-Intensity Light. Science (80–) 144:1240–1241. doi:10.1126/science.144.3623.1240

Hsu PY, Devisetty UK, Harmer SL. 2013. Accurate timekeeping is controlled by a cycling activator in Arabidopsis. Elife 2:e00473. doi:10.7554/eLife.00473

Hua Z, Vierstra RD. 2011. The cullin-RING ubiquitin-protein ligases. Annu Rev Plant Biol 62:299–334. doi:10.1146/annurev-arplant-042809-112256

Huang H, Alvarez S, Bindbeutel R, Shen Z, Naldrett MJ, Evans BS, Briggs SP, Hicks LM, Kay SA, Nusinow DA. 2016a. Identification of Evening Complex Associated Proteins in Arabidopsis by Affinity Purification and Mass Spectrometry. Mol Cell Proteomics 15:201–17. doi:10.1074/mcp.M115.054064

Huang H, Alvarez S, Nusinow DA. 2016b. Data on the identification of protein interactors with the Evening Complex and PCH1 in Arabidopsis using tandem affinity purification and mass spectrometry (TAP-MS). Data Br 8:56–60. doi:10.1016/j.dib.2016.05.014

Hurley JM, Loros JJ, Dunlap JC. 2016. Circadian Oscillators: Around the Transcription– Translation Feedback Loop and on to Output. Trends Biochem Sci 41:834–846. doi:10.1016/j.tibs.2016.07.009

Hwang JH, Seo DH, Kang BG, Kwak JM, Kim WT. 2015. Suppression of Arabidopsis AtPUB30 resulted in increased tolerance to salt stress during germination. Plant Cell Rep 34:277–89. doi:10.1007/s00299-014-1706-4

Imaizumi T, Schultz TF, Harmon FG, Ho LA, Kay SA. 2005. FKF1 F-box protein mediates cyclic degradation of a repressor of CONSTANS in Arabidopsis. Science 309:293–7. doi:10.1126/science.1110586

Imaizumi T, Tran HG, Swartz TE, Briggs WR, Kay SA. 2003. FKF1 is essential for photoperiodic-specific light signalling in Arabidopsis. Nature 426:302–6. doi:10.1038/nature02090

Ingle RA, Roden LC. 2014. Circadian regulation of plant immunity to pathogens. Methods Mol Biol 1158:273–83. doi:10.1007/978-1-4939-0700-7_18

Ito S, Song YH, Imaizumi T. 2012. LOV domain-containing F-box proteins: light-dependent protein degradation modules in Arabidopsis. Mol Plant 5:573–82. doi:10.1093/mp/sss013

Jia T, Zhang B, You C, Zhang Y, Zeng L, Li S, Johnson KCM, Yu B, Li X, Chen X. 2017. The Arabidopsis MOS4-Associated Complex Promotes MicroRNA Biogenesis and Precursor Messenger RNA Splicing. Plant Cell 29:2626–2643. doi:10.1105/tpc.17.00370

Jones MA, Williams BA, McNicol J, Simpson CG, Brown JWS, Harmer SL. 2012. Mutation of *Arabidopsis SPLICEOSOMAL TIMEKEEPER LOCUS1* Causes Circadian Clock Defects. Plant Cell 24:4066–4082. doi:10.1105/tpc.112.104828

Kevei E, Gyula P, Hall A, Kozma-Bognár L, Kim W-Y, Eriksson ME, Toth R, Hanano S, Feher B, Southern MM, Bastow RM, Viczian A, Hibberd V, Davis SJ, Somers DE, Nagy F, Millar AJ. 2006. Forward genetic analysis of the circadian clock separates the multiple functions of ZEITLUPE. Plant Physiol 140:933–45. doi:10.1104/pp.105.074864

Kiba T, Henriques R, Sakakibara H, Chua N-H. 2007. Targeted Degradation of PSEUDORESPONSE REGULATOR5 by an SCFZTL Complex Regulates Clock Function and Photomorphogenesis in Arabidopsis thaliana. PLANT CELL ONLINE 19:2516–2530. doi:10.1105/tpc.107.053033

Kikis EA, Khanna R, Quail PH. 2005. ELF4 is a phytochrome-regulated component of a negative-feedback loop involving the central oscillator components CCA1 and LHY. Plant J 44:300–313. doi:10.1111/j.1365-313X.2005.02531.x

Kim W-Y, Ali Z, Park HJ, Park SJ, Cha J-Y, Perez-Hormaeche J, Quintero FJ, Shin G, Kim MR, Qiang Z, Ning L, Park HC, Lee SY, Bressan RA, Pardo JM, Bohnert HJ, Yun D-J. 2013. Release of SOS2 kinase from sequestration with GIGANTEA determines salt tolerance in Arabidopsis. Nat Commun 4:1352. doi:10.1038/ncomms2357

Kishi T, Yamao F. 1998. An essential function of Grr1 for the degradation of Cln2 is to act as a binding core that links Cln2 to Skp1. J Cell Sci 111 (Pt 24):3655–61.

Klepikova A V., Kasianov AS, Gerasimov ES, Logacheva MD, Penin AA. 2016. A high resolution map of the *Arabidopsis thaliana* developmental transcriptome based on RNA-seq profiling. Plant J 88:1058–1070. doi:10.1111/tpj.13312

Knowles A, Koh K, Wu J-T, Chien C-T, Chamovitz DA, Blau J. 2009. The COP9 signalosome is required for light-dependent timeless degradation and Drosophila clock resetting. J Neurosci 29:1152–62. doi:10.1523/JNEUROSCI.0429-08.2009

Ko HW, Jiang J, Edery I. 2002. Role for Slimb in the degradation of Drosophila Period protein phosphorylated by Doubletime. Nature 420:673–8. doi:10.1038/nature01272

Kuroda H, Takahashi N, Shimada H, Seki M, Shinozaki K, Matsui M. 2002. Classification and expression analysis of Arabidopsis F-box-containing protein genes. Plant Cell Physiol 43:1073–85.

Larrondo LF, Olivares-Yañez C, Baker CL, Loros JJ, Dunlap JC. 2015. Circadian rhythms. Decoupling circadian clock protein turnover from circadian period determination. Science 347:1257277. doi:10.1126/science.1257277

Latres E, Chiaur DS, Pagano M. 1999. The human F box protein β-Trcp associates with the Cul1/Skp1 complex and regulates the stability of β-catenin. Oncogene 18:849–854. doi:10.1038/sj.onc.1202653

Lechner E, Achard P, Vansiri A, Potuschak T, Genschik P. 2006. F-box proteins everywhere. Curr Opin Plant Biol 9:631–8. doi:10.1016/j.pbi.2006.09.003

Lee C-M, Feke A, Li M-W, Adamchek C, Webb K, Pruneda-Paz J, Bennett EJ, Kay SA, Gendron JM. 2018. Decoys Untangle Complicated Redundancy and Reveal Targets of Circadian Clock F-Box Proteins. Plant Physiol 177:1170–1186. doi:10.1104/pp.18.00331

Lee C-M, Thomashow MF. 2012. Photoperiodic regulation of the C-repeat binding factor (CBF) cold acclimation pathway and freezing tolerance in Arabidopsis thaliana. Proc Natl Acad Sci U S A 109:15054–9. doi:10.1073/pnas.1211295109

Lee HG, Seo PJ. 2018. Dependence and independence of the root clock on the shoot clock in Arabidopsis. Genes Genomics 40:1063–1068. doi:10.1007/s13258-018-0710-4

Li S, Liu K, Zhou B, Li M, Zhang S, Zeng L, Zhang C, Yu B. 2018. MAC3A and MAC3B, Two Core Subunits of the MOS4-Associated Complex, Positively Influence miRNA Biogenesis. Plant Cell 30:481–494. doi:10.1105/tpc.17.00953

Li W, Ahn I-P, Ning Y, Park C-H, Zeng L, Whitehill JGA, Lu H, Zhao Q, Ding B, Xie Q, Zhou J-M, Dai L, Wang G-L. 2012. The U-Box/ARM E3 ligase PUB13 regulates cell death, defense, and flowering time in Arabidopsis. Plant Physiol 159:239–50. doi:10.1104/pp.111.192617

Liu J, Zou X, Gotoh T, Brown AM, Jiang L, Wisdom EL, Kim JK, Finkielstein C V. 2018. Distinct control of PERIOD2 degradation and circadian rhythms by the oncoprotein and ubiquitin ligase MDM2. Sci Signal 11:eaau0715. doi:10.1126/scisignal.aau0715

Liu T, Carlsson J, Takeuchi T, Newton L, Farré EM. 2013. Direct regulation of abiotic responses by the Arabidopsis circadian clock component PRR7. Plant J 76:n/a-n/a. doi:10.1111/tpj.12276

Lu Q, Tang X, Tian G, Wang F, Liu K, Nguyen V, Kohalmi SE, Keller WA, Tsang EWT, Harada JJ, Rothstein SJ, Cui Y. 2010. Arabidopsis homolog of the yeast TREX-2 mRNA export complex: components and anchoring nucleoporin. Plant J 61:259–70. doi:10.1111/j.1365-313X.2009.04048.x

Martin-Tryon EL, Kreps JA, Harmer SL. 2006. GIGANTEA Acts in Blue Light Signaling and Has Biochemically Separable Roles in Circadian Clock and Flowering Time Regulation. PLANT Physiol 143:473–486. doi:10.1104/pp.106.088757

Más P, Kim W-Y, Somers DE, Kay SA. 2003. Targeted degradation of TOC1 by ZTL modulates circadian function in Arabidopsis thaliana. Nature 426:567–570. doi:10.1038/nature02163

McClung CR. 2014. Wheels within wheels: new transcriptional feedback loops in the Arabidopsis circadian clock. F1000Prime Rep 6:2. doi:10.12703/P6-2

Mizoguchi T, Wheatley K, Hanzawa Y, Wright L, Mizoguchi M, Song HR, Carré IA, Coupland G. 2002. LHY and CCA1 are partially redundant genes required to maintain circadian rhythms in Arabidopsis. Dev Cell 2:629–41.

Mockler TC, Michael TP, Priest HD, Shen R, Sullivan CM, Givan SA, McEntee C, Kay SA, Chory J. 2007. The Diurnal Project: Diurnal and Circadian Expression Profiling, Model-based Pattern Matching, and Promoter Analysis. Cold Spring Harb Symp Quant Biol 72:353–363. doi:10.1101/sqb.2007.72.006

Monaghan J, Xu F, Gao M, Zhao Q, Palma K, Long C, Chen S, Zhang Y, Li X. 2009. Two Prp19– like U-box proteins in the MOS4-associated complex play redundant roles in plant innate immunity. PLoSPathog 5:e1000526. doi:10.1371/journal.ppat.1000526

Moore A, Zielinski T, Millar AJ. 2014. Online period estimation and determination of rhythmicity in circadian data, using the BioDare data infrastructure. Methods Mol Biol 1158:13–44. doi:10.1007/978-1-4939-0700-7_2

Mwimba M, Karapetyan S, Liu L, Marques J, McGinnis EM, Buchler NE, Dong X. 2018. Daily humidity oscillation regulates the circadian clock to influence plant physiology. Nat Commun 9:4290. doi:10.1038/s41467-018-06692-2

Nakamichi N, Kita M, Ito S, Sato E, Yamashino T, Mizuno T. 2005a. The Arabidopsis Pseudo– response Regulators, PRR5 and PRR7, Coordinately Play Essential Roles for Circadian Clock Function. Plant Cell Physiol 46:609–619. doi:10.1093/pcp/pci061

Nakamichi N, Kita M, Ito S, Yamashino T, Mizuno T. 2005b. PSEUDO-RESPONSE REGULATORS, PRR9, PRR7 and PRR5, Together Play Essential Roles Close to the Circadian Clock of Arabidopsis thaliana. Plant Cell Physiol 46:686–698. doi:10.1093/pcp/pci086

Nakamichi N, Kusano M, Fukushima A, Kita M, Ito S, Yamashino T, Saito K, Sakakibara H, Mizuno T. 2009. Transcript Profiling of an Arabidopsis PSEUDO RESPONSE REGULATOR Arrhythmic Triple Mutant Reveals a Role for the Circadian Clock in Cold Stress Response. Plant Cell Physiol 50:447–462. doi:10.1093/pcp/pcp004

Nakamichi N, Takao S, Kudo T, Kiba T, Wang Y, Kinoshita T, Sakakibara H. 2016. Improvement of Arabidopsis Biomass and Cold, Drought and Salinity Stress Tolerance by Modified Circadian Clock-Associated PSEUDO-RESPONSE REGULATORs. Plant Cell Physiol 57:1085–1097. doi:10.1093/pcp/pcw057

Navarro-Quezada A, Schumann N, Quint M. 2013. Plant F-Box Protein Evolution Is Determined by Lineage-Specific Timing of Major Gene Family Expansion Waves. PLoS One 8:e68672. doi:10.1371/journal.pone.0068672

Nelson DC, Lasswell J, Rogg LE, Cohen MA, Bartel B. 2000. FKF1, a clock-controlled gene that regulates the transition to flowering in Arabidopsis. Cell 101:331–40.

Ode KL, Ukai H, Susaki EA, Narumi R, Matsumoto K, Hara J, Koide N, Abe T, Kanemaki MT, Kiyonari H, Ueda HR. 2017. Knockout-Rescue Embryonic Stem Cell-Derived Mouse Reveals Circadian-Period Control by Quality and Quantity of CRY1. Mol Cell 65:176–190. doi:10.1016/J.MOLCEL.2016.11.022

Ohi MD, Vander Kooi CW, Rosenberg JA, Ren L, Hirsch JP, Chazin WJ, Walz T, Gould KL. 2005. Structural and functional analysis of essential pre-mRNA splicing factor Prp19p. Mol Cell Biol 25:451–60. doi:10.1128/MCB.25.1.451-460.2005

Onai K, Ishiura M. 2005. PHYTOCLOCK 1 encoding a novel GARP protein essential for the Arabidopsis circadian clock. Genes to Cells 10:963–972. doi:10.1111/j.1365-2443.2005.00892.x

Park BS, Eo HJ, Jang I-C, Kang H-G, Song JT, Seo HS. 2010. Ubiquitination of LHY by SINAT5 regulates flowering time and is inhibited by DET1. Biochem Biophys Res Commun 398:242–246. doi:10.1016/J.BBRC.2010.06.067

Park M-J, Seo PJ, Park C-M. 2012. CCA1 alternative splicing as a way of linking the circadian clock to temperature response in Arabidopsis. Plant Signal Behav 7:1194–1196. doi:10.4161/psb.21300

Perkins DN, Pappin DJ, Creasy DM, Cottrell JS. 1999. Probability-based protein identification by searching sequence databases using mass spectrometry data. Electrophoresis 20:3551–67. doi:10.1002/(SICI)1522-2683(19991201)20:18<3551::AID-ELPS3551>3.0.CO;2-2

Pruneda-Paz JL, Breton G, Nagel DH, Kang SE, Bonaldi K, Doherty CJ, Ravelo S, Galli M, Ecker JR, Kay SA. 2014. A Genome-Scale Resource for the Functional Characterization of Arabidopsis Transcription Factors. Cell Rep 8:622–632. doi:10.1016/j.celrep.2014.06.033

Pruneda-Paz JL, Breton G, Para A, Kay SA. 2009. A Functional Genomics Approach Reveals CHE as a Component of the Arabidopsis Circadian Clock. Science (80–) 323:1481–1485. doi:10.1126/science.1167206

Rawat R, Takahashi N, Hsu PY, Jones MA, Schwartz J, Salemi MR, Phinney BS, Harmer SL. 2011. REVEILLE8 and PSEUDO-REPONSE REGULATOR5 Form a Negative Feedback Loop within the Arabidopsis Circadian Clock. PLoS Genet 7:e1001350. doi:10.1371/journal.pgen.1001350

Reischl S, Vanselow K, Westermark PO, Thierfelder N, Maier B, Herzel H, Kramer A. 2007. β– TrCP1-Mediated Degradation of PERIOD2 Is Essential for Circadian Dynamics. J Biol Rhythms 22:375–386. doi:10.1177/0748730407303926

Risseeuw EP, Daskalchuk TE, Banks TW, Liu E, Cotelesage J, Hellmann H, Estelle M, Somers DE, Crosby WL. 2003. Protein interaction analysis of SCF ubiquitin E3 ligase subunits from Arabidopsis. Plant J 34:753–67.

Ronald J, Davis SJ. 2017. Making the clock tick: the transcriptional landscape of the plant circadian clock. F1000Research 6:951. doi:10.12688/f1000research.11319.1

Sanchez SE, Petrillo E, Beckwith EJ, Zhang X, Rugnone ML, Hernando CE, Cuevas JC, Godoy Herz MA, Depetris-Chauvin A, Simpson CG, Brown JWS, Cerdan PD, Borevitz JO, Mas P, Ceriani MF, Kornblihtt AR, Yanovsky MJ. 2010. A methyl transferase links the circadian clock to the regulation of alternative splicing. Nature 468:112–116. doi:10.1038/nature09470

Schaffer R, Ramsay N, Samach A, Corden S, Putterill J, Carré IA, Coupland G. 1998. The late elongated hypocotyl mutation of Arabidopsis disrupts circadian rhythms and the photoperiodic control of flowering. Cell 93:1219–29.

Schneider CA, Rasband WS, Eliceiri KW. 2012. NIH Image to ImageJ: 25 years of image analysis. Nat Methods 9:671–5.

Schultz TF, Kiyosue T, Yanovsky M, Wada M, Kay SA. 2001. A role for LKP2 in the circadian clock of Arabidopsis. Plant Cell 13:2659–70.

Shimizu H, Araki T, Endo M. 2015. Photoperiod sensitivity of the *Arabidopsis* circadian clock is tissue-specific. Plant Signal Behav 10:e1010933. doi:10.1080/15592324.2015.1010933

Shirogane T, Jin J, Ang XL, Harper JW. 2005. SCFbeta-TRCP controls clock-dependent transcription via casein kinase 1-dependent degradation of the mammalian period-1 (Per1) protein. J Biol Chem 280:26863–72. doi:10.1074/jbc.M502862200

Simpson CG, Fuller J, Calixto CPG, McNicol J, Booth C, Brown JWS, Staiger D. 2016. Monitoring Alternative Splicing Changes in Arabidopsis Circadian Clock GenesMethods in Molecular Biology (Clifton, N.J.). pp. 119–132. doi:10.1007/978-1-4939-3356-3_11

Somers DE, Kim W-Y, Geng R. 2004. The F-Box Protein ZEITLUPE Confers Dosage– Dependent Control on the Circadian Clock, Photomorphogenesis, and Flowering Time. PLANT CELL ONLINE 16:769–782. doi:10.1105/tpc.016808

Somers DE, Schultz TF, Milnamow M, Kay SA. 2000. ZEITLUPE Encodes a Novel Clock– Associated PAS Protein from Arabidopsis. Cell 101. doi:10.1016/S0092-8674(00)80841-7

Takase T, Ishikawa H, Murakami H, Kikuchi J, Sato-Nara K, Suzuki H. 2011. The Circadian Clock Modulates Water Dynamics and Aquaporin Expression in Arabidopsis Roots. Plant Cell Physiol 52:373–383. doi:10.1093/pcp/pcq198

Vierstra RD. 2009. The ubiquitin-26S proteasome system at the nexus of plant biology. Nat Rev Mol Cell Biol 10:385–397. doi:10.1038/nrm2688

Wang X, Wu F, Xie Q, Wang H, Wang Y, Yue Y, Gahura O, Ma S, Liu L, Cao Y, Jiao Y, Puta F, McClung CR, Xu X, Ma L. 2012. SKIP is a component of the spliceosome linking alternative splicing and the circadian clock in Arabidopsis. Plant Cell 24:3278–95. doi:10.1105/tpc.112.100081

Wang ZY, Tobin EM. 1998. Constitutive expression of the CIRCADIAN CLOCK ASSOCIATED 1 (CCA1) gene disrupts circadian rhythms and suppresses its own expression. Cell 93:1207–17.

Winter D, Vinegar B, Nahal H, Ammar R, Wilson G V, Provart NJ. 2007. An “Electronic Fluorescent Pictograph” browser for exploring and analyzing large-scale biological data sets. PLoS One 2:e718. doi:10.1371/journal.pone.0000718

Wu J-F, Tsai H-L, Joanito I, Wu Y-C, Chang C-W, Li Y-H, Wang Y, Hong JC, Chu J-W, Hsu C-P, Wu S-H. 2016. LWD-TCP complex activates the morning gene CCA1 in Arabidopsis. Nat Commun 7:13181. doi:10.1038/ncomms13181

Xie Q, Wang P, Liu X, Yuan L, Wang L, Zhang C, Li Y, Xing H, Zhi L, Yue Z, Zhao C, McClung CR, Xu X. 2014. LNK1 and LNK2 Are Transcriptional Coactivators in the *Arabidopsis* Circadian Oscillator. Plant Cell 26:2843–2857. doi:10.1105/tpc.114.126573

Xu G, Ma H, Nei M, Kong H. 2009. Evolution of F-box genes in plants: Different modes of sequence divergence and their relationships with functional diversification. Proc Natl Acad Sci 106:835–840. doi:10.1073/pnas.0812043106

Yee D, Goring DR. 2009. The diversity of plant U-box E3 ubiquitin ligases: from upstream activators to downstream target substrates. J Exp Bot 60:1109–1121. doi:10.1093/jxb/ern369

Zhang M, Zhao J, Li L, Gao Y, Zhao L, Patil SB, Fang J, Zhang W, Yang Y, Li M, Li X. 2017. The *Arabidopsis* U-box E3 ubiquitin ligase PUB30 negatively regulates salt tolerance by facilitating BRI1 kinase inhibitor 1 (BKI1) degradation. Plant Cell Environ 40:2831–2843. doi:10.1111/pce.13064

Zhou J, Lu D, Xu G, Finlayson SA, He P, Shan L. 2015. The dominant negative ARM domain uncovers multiple functions of PUB13 in Arabidopsis immunity, flowering, and senescence. J Exp Bot 66:3353–3366. doi:10.1093/jxb/erv148

Zhou Z, Wang Y, Cai G, He Q. 2012. Neurospora COP9 Signalosome Integrity Plays Major Roles for Hyphal Growth, Conidial Development, and Circadian Function. PLoS Genet 8:e1002712. doi:10.1371/journal.pgen.1002712

Zielinski T, Moore AM, Troup E, Halliday KJ, Millar AJ. 2014. Strengths and limitations of period estimation methods for circadian data. PLoS One 9:e96462. doi:10.1371/journal.pone.0096462

